# An eco-evolutionary approach to defining wildfire regimes

**DOI:** 10.64898/2026.03.17.712312

**Authors:** Sandy P. Harrison, Yicheng Shen, Olivia Haas, David Sandoval Calle, Dharma Sapkota, Iain Colin Prentice

## Abstract

Fuel availability and fuel dryness are consistently shown to be the primary drivers of wildfire intensity and burnt area. Here we hypothesise that differences in the timing of fuel build up and drying determine the optimal time for wildfire occurrence. We use gross primary production (GPP) as a measure of biomass production and hence fuel availability, and vapour pressure deficit (VPD) as a measure of fuel drying. We use the phase difference in the seasonal time course and magnitude of GPP and VPD to cluster regions that should therefore have distinct wildfire behaviour. We then show that each of the resultant clusters is distinctive in terms of one or more fire properties, specifically number of ignitions, burnt area, size, speed, duration, intensity, and length of the wildfire season. The emergence of distinct regimes as a function of two biophysical drivers reflects the fact that both vegetation and wildfire properties are a consequence of eco-evolutionary adaptions to environmental conditions. We then examine the degree to which human activities or vegetation properties modify these fire regimes within each of these clusters. Variability in GPP and VPD largely explains the within-cluster variation in fire properties. The type of vegetation cover has an influence on burnt area and carbon emissions in particular, while human activities are more important for fire properties such as size, rate of spread and duration largely through their influence of landscape fragmentation. Although both human activities and vegetation properties modify wildfire regimes, the ability to distinguish wildfire regimes using GPP and VPD alone emphasizes that land management, fire use and fire suppression are constrained by environmental conditions. This eco-evolutionary optimality approach to characterising wildfire regimes provides a basis for designing a simple fire model for Earth System modelling.

## 1. Introduction

Wildfires are ubiquitous, although their characteristics vary greatly with environmental conditions. Between 2.6±0.3% (GFED4: Giglio et al., 2013) and 5.9±0.5% (GFED5: Chen et al., 2023) of the global land area is burnt every year. Wildfires are the most important type of vegetation disturbance, necessary for the maintenance of many ecosystems, and important for initiating succession in others (Pausas and Keeley, 2009; Pausas et al., 2017; Harrison et al., 2021b; Pausas and Keeley, 2023). Wildfire-induced changes in vegetation and landscape properties also have feedbacks to climate through modulating water- and energy-exchanges and the carbon cycle (Harrison et al., 2018). Thus, an understanding of the controls on wildfire regimes is important from an Earth System perspective. Ongoing and future climate change provide an additional impetus for improved understanding of wildfire regimes, given the increased risk of more and more extreme wildfires (UNEP, 2022) and the need to mitigate this through improved management practices (Oliveras Menor et al., 2025).

Wildfire occurrence is influenced by ignitions, climate conditions, and vegetation properties, all of which may be modulated by human activity (Harrison et al., 2025). The predominant conceptual framework (Bradstock, 2010) suggests that there are four conditions, or switches, that must be met for a fire to occur: (1) there must be sufficient fuel; (2) it must be dry enough to be flammable; (3) the current weather conditions must be suitable; and (4) there must be an ignition source. Climate, vegetation, and human activity have the potential to influence each of these four switches, both independently and through their interaction.

The incidence of wildfires is highest at intermediate levels of vegetation productivity (Pausas and Bradstock, 2007; Pausas and Ribeiro, 2013; Pausas et al., 2017). This is an emergent property that reflects vegetation-climate interactions (Harrison et al., 2010; Prentice et al., 2011; Pausas and Ribeiro, 2013; Bistinas et al., 2014; Haas et al., 2022). Moisture availability is a necessary precondition for vegetation growth. Primary production is therefore enhanced as the climate becomes wetter. In general, increasing primary production increases fuel availability and thus increases the likelihood of burning. However, high levels of primary production can only be maintained when water availability is high, and thus, at the highest levels of productivity, the fuel is generally too wet to burn. This emergent relationship gives rise to two domains where fire is inhibited (Figure 1): at the dry end of the spectrum wildfires are fuel-limited, while at the wet end wildfires are dryness-limited (Harrison et al., 2010; Boer et al., 2021).

**Figure 1.**
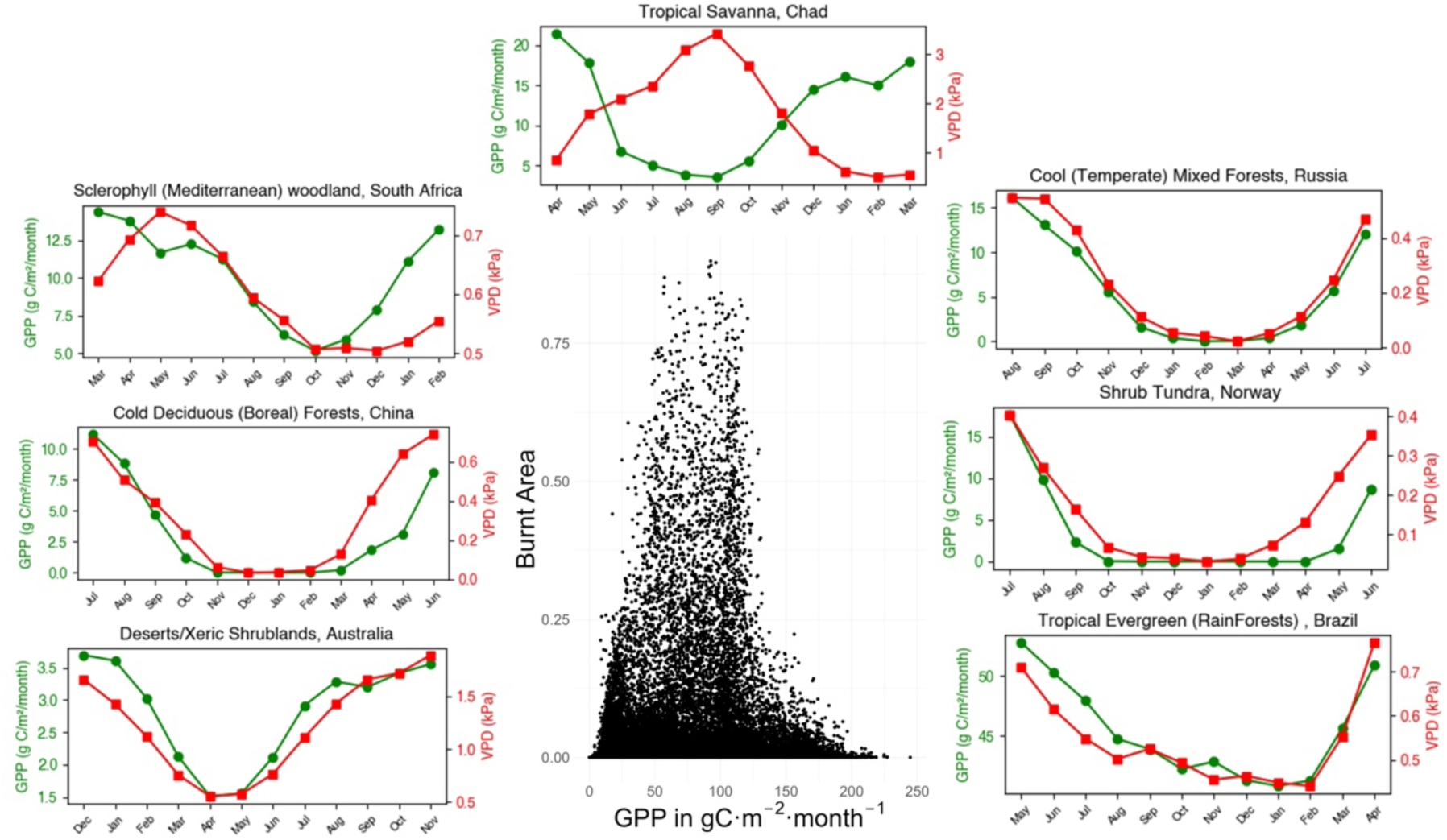
Illustration of the conceptual relationship between vegetation productivity and moisture availability, and how this influences burnt area. The middle panel illustrates the intermediate productivity hypothesis: burnt area peaks at intermediate levels of gross primary production (GPP). The surrounding panels provide examples of the observed seasonal cycle of GPP and vapour pressure deficit (VPD) at sites across different biomes. The climatological seasonal cycle was derived from observations over the interval 2003 to 2020. The panels on the left represent biomes (sclerophyll woodland, cold deciduous forest, xeric shrublands) where burnt area is largely limited by fuel availability, although impacted by differences in timing of peak GPP relative to VPD. The plots are arranged from top to bottom in order of maximum GPP. The panels on the right represent biomes (cool temperate mixed forest, shrub tundra, tropical evergreen rainforest) where burnt area is largely limited by wet conditions, although impacted by differences in timing of peak GPP relative to VPD. The plots are arranged from bottom to top in order of maximum GPP. The top panel shows an example from tropical savanna, where the maximal phasing of the seasonal cycle of GPP and VPD produces conditions that are optimal for burning.

The intermediate productivity hypothesis is consistent with the spatial patterns in observed fire occurrence. Boer et al. (2016, 2021), for example, have shown that these patterns in the probability of wildfire occurrence can be determined as a function of annual precipitation and annual evapotranspiration. In this context annual precipitation is a control on fuel availability while annual evapotranspiration is a measure of fuel drying. However, the intermediate productivity hypothesis does not provide an immediate explanation for the temporal patterns of fire occurrence and the timing of the fire season, although these should also reflect the interplay between fuel availability and fuel dryness. A number of studies have recognised the importance of rainfall seasonality in controlling the occurrence and timing of wildfires (Bowman et al., 2009; Bowman et al., 2014; Ellis et al., 2023). Antecedent conditions that determine fuel build-up have been shown to play an important role in the timing of fire occurrence. In fuel-limited regions, burnt area has been shown to be sensitive to vegetation productivity in the several months before the fire through influencing fuel build-up, but that current weather conditions which influence fuel drying are more important in dryness-limited regions (Kuhn-Régnier et al., 2021; Haas et al., 2024). Similarly, burnt area is also sensitive to anomalously dry conditions prior to the fire, as indexed by accumulated number of dry-days, which limits fuel-build up and therefore reduces burnt area (Kuhn-Régnier et al., 2021; Haas et al., 2024). These findings suggest that differences in the phasing of the controls on fuel build-up and drying may be crucial in explaining fire occurrence.

Wildfires are often described in terms of so-called fire regimes, where a regime represents a particular combination of fire characteristics, such as ignition source, type (ground, crown), seasonality, frequency, intensity, size or extent (Bond & Keeley, 2005; Lavorel et al., 2007; Kelly et al., 2023). Thus, the concept of a fire regime seeks to link both the spatial and temporal characteristics of wildfires. In general, such definitions adopt a top-down perspective to delineate specific wildfire regimes. However, there have been a number of studies that have characterised wildfire regimes more objectively based on different wildfire properties. Archibald et al. (2013), for example, used a Bayesian clustering approach to defining fire regimes, or “pyromes” based on remote-sensing data on fire size, frequency, intensity, season and extent. They identified five pyromes, four characterising natural wildfire regimes and the fifth delineating wildfires influenced by human activities. Garcia et al. (2022) identified four regimes based on remotely-sensed burned area and active fire records for the period 2001 to 2020 using k-means clustering. In the most comprehensive analysis to date, Pais et al (2023) used several clustering and machine-learning approaches to identify wildfire regimes using remotely sensed fire data from 2000-2018, identifying 15 different pyromes which they further subdivided into 62 distinct regional regimes. Although each of these studies considers seasonality, they do not consider what drives differences between the regimes and thus regions with low fire, such as deserts and tropical forests, are grouped together regardless of whether the absence of fire is a result of fuel or dryness.

There are other problems with the classification of wildfire regimes using observed wildfire properties. First, wildfire occurrence is highly stochastic and this poses problems for identifying regimes using observations of wildfire properties over recent decades, especially in regimes with infrequent fires such as boreal forests. Second, classifications based on recent observations make the assumption that wildfire properties are strongly coupled and so some combinations of wildfire properties cannot exist in the real world. However, model simulations of wildfires under past and future climates indicate that this is not the case, and that there can be a decoupling between, for example, wildfire occurrence and intensity (Haas et al., 2023; Haas et al., 2025). The focus on emergent properties as opposed to the fundamental drivers of wildfire is also problematic. The biophysics linking key drivers to wildfire occurrence does not change under different climate states, whereas the emergent properties could change. Current patterns in the magnitude and seasonal cycle of GPP reflect modern climate gradients. However, atmospheric CO_2_ concentration also impacts GPP through changing water-use efficiency (Ainsworth & Rogers, 2007; Keenan et al., 2013; Prentice et al., 2014). Increases or decreases in atmospheric CO_2_ concentration thus have an impact on GPP patterns independent of climate.

An alternative approach is to use fire drivers to define regimes and to determine the wildfire properties associated with them (e.g. Zubkova et al., 2022). Eco-evolutionary optimality (EEO) theory (Franklin et al., 2020; Harrison et al., 2021a), which is predicated on the assumption that plants are adapted to the environments in which they occur, has been shown to provide a useful and sometimes surprisingly simple approach to understanding vegetation properties from the leaf-level coupling between photosynthesis and stomatal conductance (Wang et al., 2017; Stocker et al., 2020) to community assembly patterns (Joshi et al., 2022). This EEO approach can be extended to fire regimes, in the sense that plants must also be adapted to the fire regimes characteristic of particular environmental conditions. It implies that, on multi-decadal or longer timescales, the environmental drivers of vegetation properties and wildfire regimes are linked.

Human actions can influence wildfires directly (via increasing or suppressing ignitions) or indirectly via landscape modification, including fragmentation of vegetation. Although humans are considered responsible for most ignitions (Farid et al., 2024), most human-set fires are for agricultural purposes and tend to be small and quickly suppressed (Jones et al., 2022; Millington et al., 2022). Deliberately set fires, fires used for land clearing, or fires that escape from agricultural land into natural vegetation, may be larger but are less common. Policies that have led to long-term wildfire suppression, however, has led to an increase in fuel loads and this has led to larger and more intense wildfires in many parts of the world given suitable climate conditions (Moreira et al., 2020; Jones et al., 2022; Kreider et al., 2025). Despite this, human modification of the landscape is a more pervasive influence on wildfires than fire suppression. Landscape fragmentation generally leads to reduced fuel continuity and hence often a reduction in wildfires (Andela et al., 2017; Jones et al., 2022; Zubkova et al., 2023). However, the impact of landscape fragmentation varies with vegetation type: in fire-adapted vegetation, fragmentation has little impact on the wildfire regime whereas in ecosystems that are characterised by infrequent fires, fragmentation creates environmental conditions that favour increased fire (Harrison et al., 2021b). The introduction of crops and grazing also modify landscape properties, but environmental factors are a strong determinant of the suitability of a region for agriculture (Ramankutty et al., 2002; Sloat et al., 2020). Thus, while human actions are a proximal cause of changes in wildfire, they are themselves subject to environmental constraints that should make it possible to incorporate them within an environmentally-driven EEO framework.

In this paper, we explore the possibility of using an EEO-based approach to defining and characterising wildfire regimes on the basis of their key drivers. We first develop a theory based on the idea that differences in the timing of fuel build up and fuel drying determine the optimal time for fire occurrence and hence the timing and length of the fire season. Then, using gross primary production (GPP) as a measure of fuel availability, and vapour pressure deficit (VPD) as a measure of fuel drying, we cluster the world into regions that should, theoretically, be associated with distinct wildfire regimes. We derive GPP using a universal EEO-based model of primary production, since this is not an observed quantity globally. We then demonstrate that these pyroclimate regions differ from one another in terms of fire properties, including frequency, intensity, the length of the fire season and burnt area. Finally, we examine the impact of differences in vegetation properties and human activities in modulating the fire regimes within each of these pyroclimate regions.

## 2. Theory

Many factors related to climate, vegetation, landscape characteristics and human activities are considered as important influences on wildfire (Bowman et al., 2009; Harrison et al., 2010; Bowman et al., 2011), but GPP and VPD have consistently been found to be the key drivers of wildfires and their properties (Haas et al., 2024). Net primary production (NPP) is the fundamental measure of fuel production but is proportional to GPP. Although this relationship varies to some extent between different vegetation types (Collati and Prentice, 2019), GPP nevertheless provides a reasonable approximation for fuel availability, while the magnitude of VPD controls the rate of fuel drying. Differences in the seasonal cycle of these two properties are important (Krawchuk and Moritz, 2011; Kuhn-Régnier et al., 2021). High GPP prior to the wildfire season indicates high fuel loads, but high GPP during the fire season can only occur if precipitation is high and this precludes fuel drying. High VPD before the fire season means that plants are likely to be droughted and fuel loads therefore correspondingly low, whereas high VPD during the fire season means that the available fuel will be sufficiently dry to burn. These relationships are consistent with the intermediate productivity hypothesis, as discussed above. Ignitions, whether natural or anthropogenic, are not considered limiting (Haas et al., 2024) in this framework.

Comparisons of the seasonal cycle of GPP and VPD at individual sites show distinct patterns across different biomes (Figure 1). Anti-phasing of GPP and VPD is characteristic of savannas and tropical grasslands, for example, whereas the seasonal cycles of GPP and VPD are in phase in tundra and in temperate and tropical evergreen forests. Muted changes in GPP and VPD are characteristic of relatively wet tropical evergreen forests. Differences between biomes in both the relative magnitude of GPP and VPD, and in the relative length of the peaks of the two variables, are also apparent. To characterise the environmental niche for wildfire in any place, we therefore use a simple clustering approach based on the seasonal cycle of GPP and VPD using a climatology for the interval 2003-2020.

## 3. Results

We identified 18 statistically distinct clusters. The geographic distribution of each cluster (Fig 2; Supplementary Figures 1-18; Table 1) makes intuitive sense in terms of broadscale climate and vegetation patterns. There are distinct latitudinal clusters in the northern extratropics that are consistent with the transitions from tundra through cold deciduous forest, to boreal forest and ultimately temperate forest. There are also distinct clusters in the tropics broadly corresponding to transitions between evergreen tropical forests and more open savanna and ultimately xeric vegetation. The chosen spatial resolution of the input data (10 km or 50 km) does not affect the number of clusters or the geographic distribution of clusters (Supplementary Figure 19). The area occupied by individual clusters (Table 1) varies by an order of magnitude: cluster 18 occupies the smallest area (2.4 Mm^2^), cluster 17 the largest area (16.7 Mm^2^).

**Figure 2.**
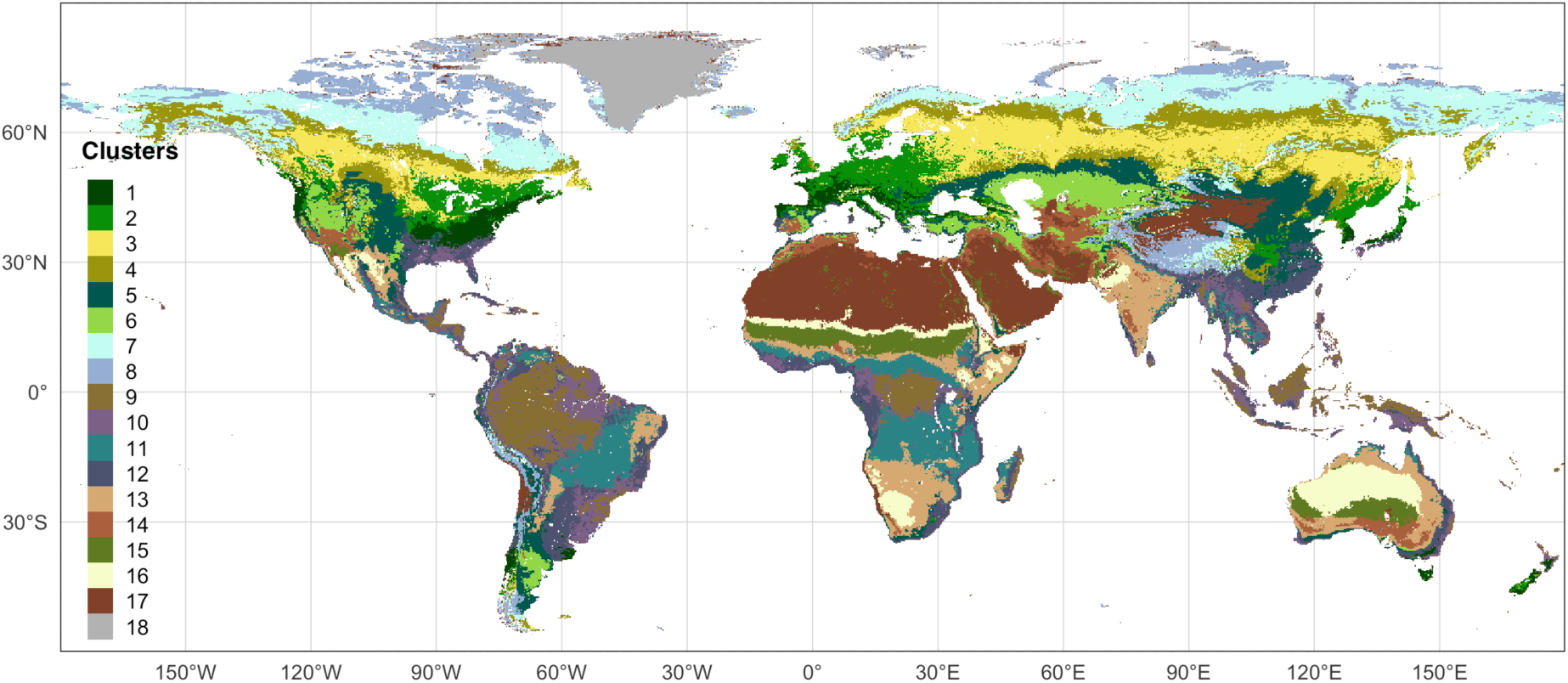
Regions with similar patterns in the magnitude and seasonal phasing of gross primary production (GPP) and vapour pressure deficit (VPD) based on X-means clustering of data at 10 km resolution. Maps showing the geographic distribution of each individual cluster are given in Supplementary Material.

**Table 1.** Area occupied by individual clusters, expressed in total area (to the nearest km^2^) and as a percentage of the total land area.

| Cluster | Total area (km<sup>2</sup>) | % Land Area |
| --- | --- | --- |
| 1 | 3033240 | 2.29 |
| 2 | 6109011 | 4.62 |
| 3 | 9166104 | 6.93 |
| 4 | 7958016 | 6.01 |
| 5 | 9223446 | 6.97 |
| 6 | 4783995 | 3.62 |
| 7 | 9248502 | 6.99 |
| 8 | 5800389 | 4.38 |
| 9 | 8157999 | 6.16 |
| 10 | 7630228 | 5.77 |
| 11 | 8674658 | 6.56 |
| 12 | 9981368 | 7.54 |
| 13 | 10255912 | 7.75 |
| 14 | 3691312 | 2.79 |
| 15 | 4467657 | 3.38 |
| 16 | 5107766 | 3.86 |
| 17 | 16651346 | 12.58 |
| 18 | 2394408 | 1.81 |

Each of the resultant clusters is distinctive in terms of one or more fire properties, and therefore represents a unique fire regime. The Hotellings *T^2^* statistic, which evaluates the hypothesis of difference between the multivariate medians of the seven fire properties, shows that only one pair of clusters (8 and 18) are statistically indistinguishable (p = <0.01) from one another. The choice of significance level was based on the modest sample size (153 potential comparisons) and the identification of a single non-significant relationship does not exceed the false negative rate. We used pairwise Wilcoxon tests to identify which fire properties distinguished the different clusters. The degree of differentiation varied between fire properties, with clusters being more distinct from one another with respect to burnt area, fire size and carbon emissions (ca 95% at p =<0.01, Table 2) than other properties. Nevertheless, there were significant differences between more than 85% of the clusters for all fire properties (Table 2, Figure 3). Some clusters were significantly different from all other clusters with respect to one or more properties (Figure 3, Supplementary Table 1). Clusters 4, 9, 10, 13 and 16, for example, were distinct from all other clusters with respect to average fire size, while Clusters 3, 4, 5, 11, 12, 14 and 15 were distinct from all other clusters with respect to burnt area fraction. Some clusters were only distinct from all other clusters with respect to a single property (cluster 2 for carbon emissions, cluster 6 for average fire speed, cluster 12 for burnt area fraction, cluster 17 for number of ignitions). However, there were only three clusters (cluster 1, cluster 7, cluster 18) that shared properties with at least one other cluster for all of the fire properties, although the cluster from which they were statistically indistinguishable varies between properties (Figure 3).

**Figure 3.**
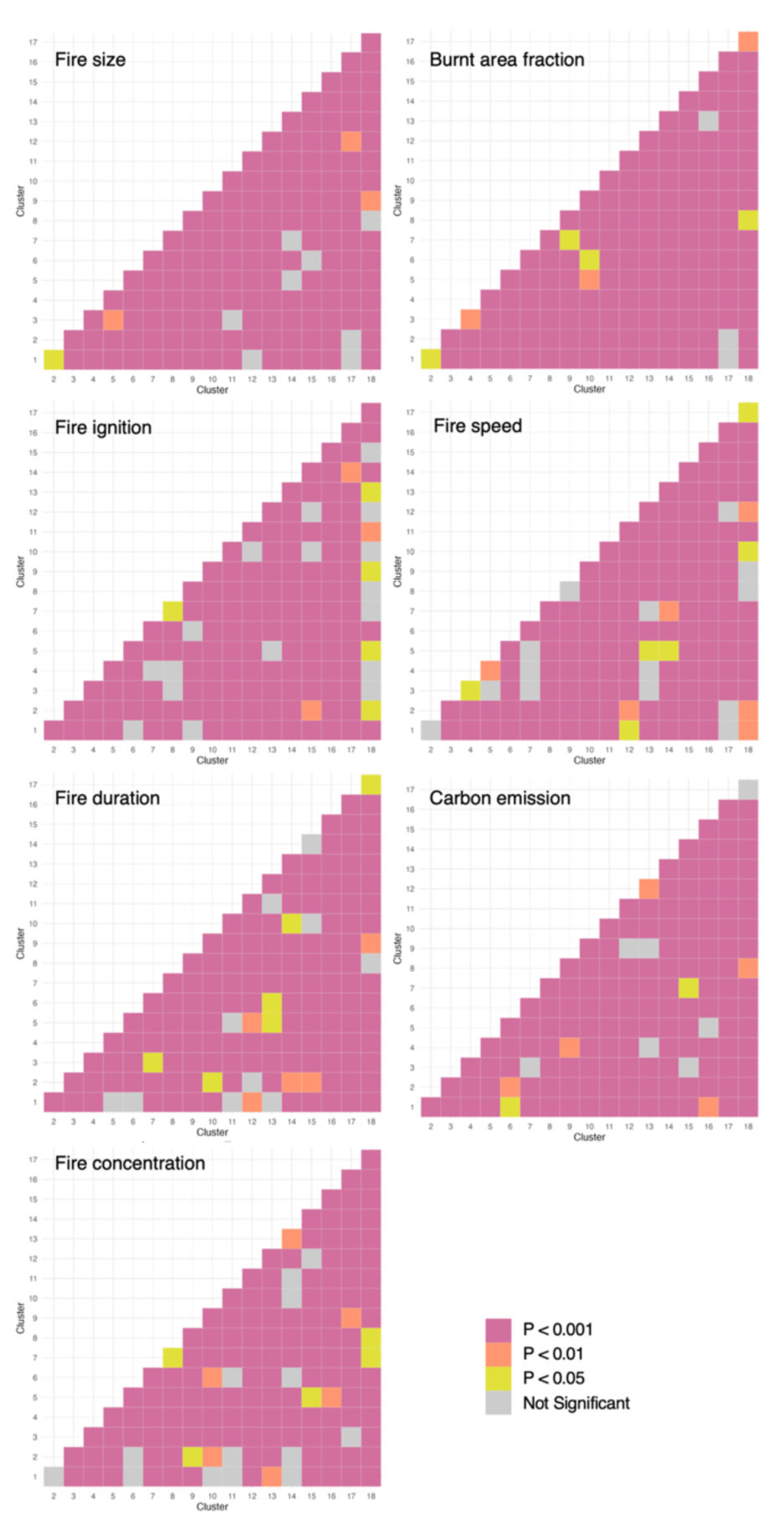
Pairwise comparison of clusters with respect to individual fire properties: average fire size (km^-2^), burnt area fraction (%), number of ignitions (km^-2^), average fire speed (km/day^-1^), average fire duration (days), carbon emission (gCm^-2^) and fire seasonal concentration (unitless). Significance was evaluated at multiple levels (p = 0.001, 0.01 and 0.05 levels) for the 153 comparisons, where differences at p < 0.01 were considered to minimise the influence of false positives.

**Table 2:** Summary of the paired Wilcoxon tests. The table shows the percentage of clusters that are significantly different from the other clusters with respect to a specific fire property at different p values (<0.001, <0.01, <0.05). The fire properties are: average fire size (km^-2^), burnt area fraction (%), number of ignitions (km^-2^), average fire speed (km/day^-1^), average fire duration (days), carbon emission (gCm^-2^) and fire seasonal concentration (unitless).

| Fire property | p <0.001 | p <0.01 | p <0.05 |
| --- | --- | --- | --- |
| Burnt area | 93.46 | 95.42 | 98.04 |
| Size | 92.16 | 94.12 | 94.77 |
| Speed | 83.01 | 86.93 | 90.85 |
| Ignitions | 83.66 | 85.62 | 88.89 |
| Duration | 86.27 | 89.54 | 93.46 |
| Emissions | 90.85 | 94.12 | 95.42 |
| Concentration | 83.66 | 87.58 | 90.85 |

The relationships between magnitude and relative timing of GPP and VPD and fire properties (Figure 4) illustrate the importance of biophysical drivers on fire regimes and is consistent with current theoretical frameworks. Burnt area and fire size tend to be higher when VPD is high and there is maximum separation between peak GPP and peak VPD, consistent with our hypothesis that the optimum condition for fires occurs when there is sufficient vegetation growth for fuel build up followed by an interval when climatic conditions allow for fuel drying. Carbon emissions are maximal at intermediate levels of GPP, consistent with the intermediate productivity hypothesis, though this relationship is less apparent for burnt area. The number of ignitions and the seasonal concentration tend to both be highest when the degree of separation between peak GPP and peak VPD is low, but show no consistent relationship with GPP although they tend to occur at intermediate levels of VPD, suggesting that these properties largely reflect agricultural fires.

**Figure 4.**
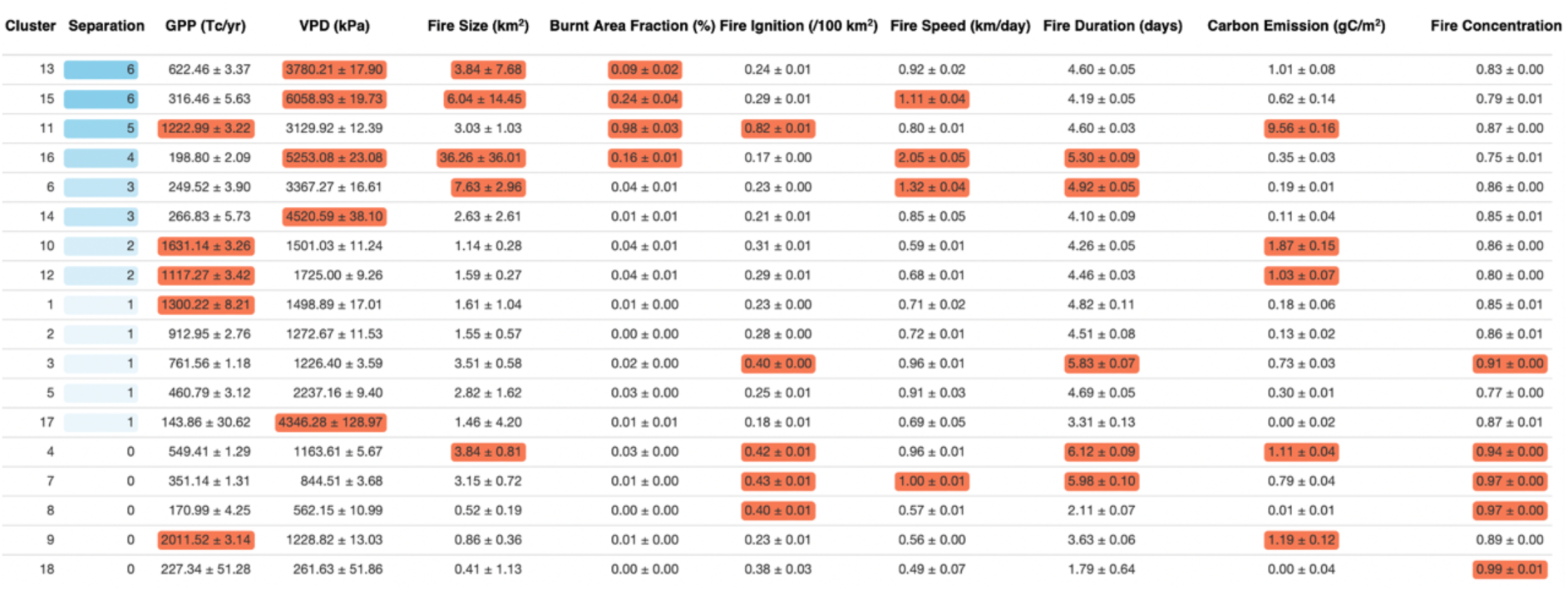
Summary of the properties of each cluster. The clusters are ordered in terms of their degree of separation of peak gross primary production (GPP) and vapour pressure deficit (VPD), expressed in number of months. The red coloured boxes indicate clusters in the top quartile for GPP, VPD, fire size, burnt area fraction, number of ignitions, fire speed, fire duration, carbon emissions and fire concentration.

Within-cluster variability in burnt area and carbon emissions is strongly controlled by GPP and VPD, with one or other being the dominant driver in ten of the twelve cases where significant relationships were found (Table 3). The GLM models which do not include these variables generally have much lower R^2^ (Supplementary Table 2). The impact of VPD on burnt area is always positive, consistent with the fact that increasing dryness will promote increased burnt area. The impact of GPP on burnt area is generally positive, consistent with the idea that increasing GPP leads to higher fuel loads. The one exception, where GPP is an important control but has a negative effect on burnt area is cluster 9, which broadly corresponds to regions dominated by tropical rainforest (Supplementary Figure 9) where higher GPP implies conditions where the fuel is likely to be too wet to burn. The impact of VPD on carbon emissions is always positive, again reflecting the importance of dry fuels for completeness of burn. The impact of GPP on carbon emissions is positive except in clusters 3 and 6. Cluster 3 broadly corresponds to the northern boreal forest (Supplementary Figure 3), and the negative impact of GPP appears to reflect the importance of vegetation cover as drivers in the model. The apparently negative impact of GPP on emissions in Cluster 6 (Supplementary Figure 6) is probably spurious given that the overall R^2^ for this model is very low (0.03).

**Table 3:** Summary of results for the generalised linear model (GLM) of the within-cluster variability in fire properties as a function of the annual gross primary production (GPP, gCyr^-1^), maximum VPD (VPD, Pa), fractional grass cover (G), fractional shrub cover (S), fractional tree cover (T), fractional cropland cover (C), fractional pasture cover (P), road density (R) and population density (Pn). R^2^ values for significant models are given in **bold**; the significant factors (F) are given in order of importance and colour-coded to indicate whether their impact on a fire property is positive (in red) or negative (in blue). Details of the individual models are given in Supplementary Tables 3-9.

|  | BA |  | Ignitions |  | Size |  | Speed |  | Duration |  | Emissions |  | Concentration |  |
| --- | --- | --- | --- | --- | --- | --- | --- | --- | --- | --- | --- | --- | --- | --- |
| cluster | R2 | F | R2 | F | R2 | F | R2 | F | R2 | F | R2 | F | R2 | F |
| 1 | **0.34** | VPD<br>GPP S | **0.04** | ns | **0.33** | VPD<br>G | **0.11** | VPD<br>C | **0.16** | VPD | **0.58** | GPP<br>VPD S<br>G | **0.07** | P R C |
| 2 | **0.30** | ns | **0.08** | R GPP<br>S | **0.27** | VPD<br>C | **0.14** | S<br>VPD | **0.06** | VPD | **0.19** | ns | **0.07** | GPP<br>VPD<br>R |
| 3 | **0.24** | VPD<br>G T P<br>R | **0.1** | VPD<br>G P T | 0.14 | ns | **0.1** | R G | **0.19** | S<br>GPP<br>VPD<br>T R | **0.26** | VPD R<br>GPP G<br>S P | **0.22** | VPD<br>T C S<br>R GPP |
| 4 | **0.29** | VPD<br>P S R<br>G | **0.11** | S P<br>GPP<br>VPD<br>G C T<br>R | 0.17 | ns | **0.12** | C<br>VPD<br>S | **0.23** | S<br>VPD<br>T C P<br>G | **0.30** | VPD S | **0.36** | C<br>VPD<br>T R G<br>Pn |
| 5 | **0.11** | C G P<br>GPP T | **0.34** | C G S | 0.32 | ns | **0.22** | C P<br>T | **0.05** | S T P | **0.22** | GPP C<br>VPD | **0.05** | GPP C<br>VPD |
| 6 | **0.31** | ns | **0.24** | C S R<br>G<br>VPD | 0.34 | ns | **0.32** | C T<br>R G<br>P | **0.14** | C S | **0.25** | GPP S<br>T P | **0.03** | GPP |
| 7 | **0.31** | VPD<br>T GPP | **0.09** | GPP S<br>VPD<br>G P R | 0.2 | ns | **0.14** | VPD<br>T G<br>R<br>GPP | **0.18** | VPD<br>T<br>GPP<br>R | **0.32** | VPD T<br>S G | **0.05** | R C<br>GPP |
| 8 | **0.22** | GPP | **0.31** | VPD R<br>T S | **0.37** | VPD<br>G | **0.04** | ns | **0.2** | GPP | **0.39** | S GPP<br>VPD T<br>R | **0.18** | C R G<br>GPP |
| 9 | **0.45** | VPD<br>GPP<br>G T | **0.24** | VPD<br>G P | **0.12** | VPD | **0.12** | VPD | **0.11** | VPD<br>R T | **0.41** | VPD T<br>S G P | **0.03** | VPD<br>T GPP |
| 10 | **0.31** | ns | **0.17** | VPD S<br>R | **0.08** | ns | **0.04** | C T | **0.02** | T | **0.28** | VPD S | **0.11** | S G T |
| 11 | **0.38** | G T R<br>VPD<br>S P | **0.37** | G T P<br>C R<br>VPD | 0.31 | ns | **0.25** | S<br>VPD<br>P R<br>C | **0.19** | P G S<br>C R<br>GPP | **0.49** | T G<br>VPD<br>GPP P<br>R S | **0.13** | T<br>VPD<br>G P |
| 12 | **0.29** | ns | **0.16** | P R<br>GPP<br>VPD S | **0.24** | VPD<br>GPP<br>S C<br>P G | **0.14** | GPP<br>VPD<br>S T<br>C P | **0.04** | S P<br>GPP | **0.28** | GPP<br>VPD S<br>P R | **0.05** | T P Pn<br>S G |
| 13 | **0.63** | GPP<br>VPD<br>S G T<br>R P | **0.43** | GPP<br>VPD P<br>G C | 0.52 | ns | **0.36** | C<br>VPD<br>P T<br>S<br>GPP | **0.16** | P C S<br>GPP<br>VPD | **0.71** | GPP<br>VPD S<br>G P R<br>T | **0.03** | VPD P |
| 14 | **0.68** | ns | **0.35** | P GPP<br>VPD S | **0.29** | S C | **0.22** | S C<br>GPP | **0.11** | S<br>GPP | **0.69** | GPP<br>VPD P | **0.03** | GPP P<br>VPD |
| 15 | **0.78** | GPP S<br>VPD<br>P C<br>Pn | **0.62** | GPP S<br>G T P | 0.46 | ns | **0.31** | C<br>GPP<br>VPD | **0.09** | VPD<br>C | **0.9** | GPP S<br>C T G<br>R | **0.06** | VPD<br>R Pn |
| 16 | 0.68 | ns | **0.09** | C P | 0.46 | ns | 0.38 | ns | **0.34** | GPP<br>G R C<br>VPD<br>S | 0.68 | ns | **0.04** | GPP<br>VPD<br>C |
| 17 | **0.53** | GPP | **0.11** | VPD | **0.2** | ns | **0.2** | VPD<br>G<br>GPP<br>C | **0.19** | GPP | **0.68** | GPP<br>VPD C<br>P S R | **0.05** | GPP<br>VPD |
| 18 | **0.73** | GPP<br>VPD<br>S | 0.52 | ns | **0.91** | ns | **0.61** | ns | 0.47 | ns | **0.73** | GPP T | **0.59** | P |

Human activities are not identified as important in most models of burnt area and carbon emission, but are more frequently identified as important in determining numbers of ignitions, fire size, speed and duration (Table 3). They are rarely the dominant factor except in the case of crop cover, which is the dominant factor for fire speed in six of the pyroclimate clusters. Crop cover always has a negative impact on fire size, speed and duration, reflecting the fact that increasing crop cover leads to increased landscape fragmentation and cropping also reduces the annual fuel load. Road density has a negative impact on fire speed and duration, also reflecting landscape fragmentation, but does not appear as an important determinant of fire size. The influence of crops and roads on number of ignitions is more nuanced: crop area always has a positive influence on number of ignitions whereas road density generally has a negative influence. Given that these factors have a negative influence on fire size, speed and duration, this suggests that the positive relationship between crop cover and number of ignitions reflects agricultural burning. This suggestion is borne out by the importance of crop cover in the models for clusters representing sub-Saharan and southern Africa (Supplementary Figure 11, 16). Road density has a negative influence of number of ignitions, except in cluster 4, which corresponds to the northern boreal forest (Supplementary Figure 4). In this case, the positive relationship likely reflects the fact that the creation of roads in an ecosystem which is not densely settled and also not adapted to fire both increases human access and creates edge effects such as increased fuel drying and increased wind speed, and hence increases the probability of fire. Population density does not emerge as a significant influence on fire properties, except in one model for burnt area and three models for seasonal concentration, but in no case is it the dominant influence. Thus, while human activities modulate fire behaviour in predictable ways, they do not give rise to substantially different fire regimes or overwhelm the predominant influence of GPP and VPD on these regimes.

Vegetation properties contribute to the variability of burnt area, number of ignitions, fire speed, fire duration and carbon emissions (Table 3), though they are rarely the most important factor. Shrub cover is the most likely of the vegetation properties to be influential, but nevertheless is only the dominant factor for fire speed in three of the clusters and for fire duration in five of the clusters. Shrub cover always has a positive effect on burnt area and fire speed but generally has a positive effect on number of ignitions, fire duration and carbon emissions. Shrub cover has a negative relationship with number of ignitions in Clusters 5 and 6 (Supplementary Figures 5 and 6), for example, and this appears to be related to the importance of crop cover in these models. The positive relationship between ignitions and crop cover points to the use of fire for agricultural purposes, in which case increasing shrub cover would correspond to non-agricultural land and have a dampening effect on ignitions. Tree and grass cover are included in some of the models: tree cover generally has a negative relationship, while grass cover generally has a positive relationship with fire properties. These relationships support the idea that extensive grass cover promotes rapidly spreading ground fires, whereas increases in tree cover will reduce the incidence of such fires.

## 4. Discussion and Conclusions

We have used the two most important drivers of wildfire behaviour, namely the seasonal phasing and relative magnitude of GPP and VPD, to identify distinct pyroclimate regions and shown that this regionalisation can be mapped on to wildfire properties. This provides a more nuanced approach to dividing up the continuum of wildfire regimes than previous attempts to categorise these regimes based on the wildfire properties themselves. The regions show distinct geographic patterns which broadly correspond to biomes, which is unsurprising given that biomes themselves are largely determined by climate (Prentice et al., 1992). The use of X-means clustering with a Bayesian information criterion to determine the boundaries between clusters removes the need to predetermine a number of clusters and identifies breaks in observed behaviour between clusters (Pelleg & Moore, 2000; Radwan et al., 2020).

Our analysis emphasises that wildfire properties can vary independently across different regimes. It is possible, for example, to have high burnt area associated with high or low numbers of ignitions and with high or low wildfire intensity. Similarly, it is possible to have highly seasonal fire regimes associated with either low or high burnt area. This has been pointed out in previous studies (e.g. Archibald et al., 2013) and is consistent with the idea that wildfire properties are decoupled and may change independently as climate shifts (Haas et al., 2023; Haas et al., 2025). The recent emergence of novel fire regimes (Moreira et al., 2020; Jones et al., 2022; Kreider et al., 2025; Shen et al., 2025) and the likelihood that there will be major changes in wildfire properties under future climate change (Haas et al., 2025; Shen et al., 2025) means that it is important to consider this potential decoupling in the ongoing development of global fire models, many of which explicitly tie some of these properties together (Rabin et al., 2017).

Previous attempts at defining wildfire regimes have identified distinct anthropogenic pyromes. Some clusters are indeed characterised by high levels of anthropogenic activity, but the relative importance of different indices of such activity varies between regions. In regions with high levels of human activity, factors associated with landscape fragmentation such as population density, road density and cropland area act to reduce wildfires overall (Knorr et al., 2014; Andela et al., 2017; Harrison et al., 2025). In contrast, the variability in pastureland has a more complex impact. Some regions have high levels of pastureland associated with low population density, and here the overall impact is to increase wildfires (Ruiz-Mirazo et al., 2012); but in more fire-prone regions high levels of pastureland leads to reduced burnt area and carbon emissions. The fact that our regionalisation is dependent only on biophysical drivers of wildfire means that none of the identified wildfire regimes can be considered as anthropogenic, although human activities modify the wildfire regimes within these clusters in predictable and consistent ways.

Our analyses are based on data covering the interval 2003 to 2020. The highly stochastic nature of wildfire occurrence means that this is a short time window for characterising fire regimes in regions where fire return times are decades to centuries. As a result, it is likely that we do not capture the full range of wildfire behaviour boreal and tundra ecosystems. Nevertheless, the methodology does distinguish several clusters in the northern extra-tropics, implying that there should be differences between these regions in fire occurrence and behaviour.

There are multiple interactions between climate, vegetation and wildfire, with vegetation properties or traits often considered important in determining wildfire regimes (Enright et al., 2014; Kramer et al., 2016; Prior et al., 2017). Eco-evolutionary theory provides a simple approach to understanding vegetation distribution and vegetation traits as being optimal adaptations to ensure ecosystem functioning under a given climate (Franklin et al., 2020; Harrison et al., 2021a). The idea that environmental factors shape vegetation patterns can also be extended to fire regimes, since species and fire regimes have co-evolved (Keeley et al., 2011; Archibald et al., 2018). Pausas et al (2025) have shown, for example, that the different incidence of crown versus ground fires in boreal forests in North America and Eurasia reflects the warmer and more productive environments in Eurasia than in North America. Temperature and moisture availability are key drivers of plant productivity, driving trade-offs between carbon assimilation and water use that underpin photosynthesis, plant growth and competition (Cai and Prentice, 2020; Cai et al., 2025). The ability to delineate wildfire regimes based on these same biophysical drivers opens up the possibility of modelling wildfires using an EEO approach. Given the current limitations of the more mechanistic fire-enable vegetation models used to predict changing wildfire regimes (Kloster and Lasslop, 2017; Hantson et al., 2020; Li et al., 2024), a simpler approach based on EEO and driven by simulated VPD and GPP may well provide a useful alternative in an Earth System modelling context.

The simple EEO-based approach to delineating pyroclimates described here is only a first step towards the development of a new wildfire model. GPP is a useful surrogate for fuel loads, particularly live fuel loads which drive both ground and crown fires in many regions of the world (Nolan et al., 2022). However, the accumulation of dead fuel over time is also a critical factor in determining the occurrence and spread of wildfires, and thus a realistic assessment of fuel availability would require information of both multi-annual accumulation and associated decomposition rates (McNorton & Di Giuseppe, 2024). Plant traits are known to affect both fuel flammability and the rate of decomposition (Grootemaat *et al*. 2015; Archibald *et al*. 2018; Dai *et al*. 2025) and will likely need to be taken into account in order to assess fuel loads and their impacts on the timing and interannual variability of wildfires. However, the plant traits that influence flammability and rates of decomposition reflect prevailing environmental conditions including the characteristic fire regime (Bond & Keeley, 2005; Jaureguiberry & Díaz 2023; Dai *et al*. 2025) and this opens up the possibility of modelling fuel loads from an EEO perspective. This would both complement and extend our current EEO framework for delineating wildfire regimes.

## 5. Methods

We used an EEO-based model, the P model (Stocker et al., 2020) to calculate the seasonal cycle of GPP. The P model is a light-use efficiency model based on the Farquhar-von Caemmerer-Berry photosynthesis model (Farquhar et al., 1980) for instantaneous biochemical processes and uses two EEO-hypotheses, the coordination (Maire et al., 2012; Wang et al., 2017) and least-cost hypotheses (Wright et al., 2003; Prentice et al., 2014; Wang et al., 2017), to account for the spatial and temporal acclimation of carboxylation and stomatal conductance to varying environmental conditions on weekly to monthly timescales. The P model has been extensively validated at site and ecosystem level in terms of basic physiological characteristics related to photosynthesis (Smith et al., 2019; Jiang et al., 2020; Peng et al., 2020; Smith and Keenan, 2020; Harrison et al., 2021) and shown to predict the geographic patterns of GPP under modern conditions successfully (Wang et al., 2017; Stocker et al., 2020) as well as the recent trends in GPP as measured at eddy-covariance flux sites (Cai and Prentice, 2020) and against remote-sensing data comparable to the best-performing dynamic global vegetation models (Cai et al., 2025).

The P model simulates GPP as a function of atmospheric CO_2_ concentration, air temperature, atmospheric pressure, vapour pressure deficit (VPD), and incident photosynthetic photon flux density (PPFD), with an empirical function to account for soil moisture stress (Stocker et al., 2020). Soil moisture limitation was estimated using version 2 of the Simple Process-Led Algorithm for Simulating Habitats (SPLASH v2.0: Sandoval et al., 2024). The climate data required to force the P model and SPLASHv2, and to calculate the seasonal cycle of VPD, were obtained from ERA5-Land (Muñoz-Sabater, 2019) at monthly timesteps. The soil data required as additional input to SPLASHv2 was derived from Soilgrids v2.0 (Poggio et al., 2021). The P model was run from 2003 to 2020, using fAPAR data from Jeong et al. (2024) and annually varying CO_2_ from the NOAA Global Monitoring Laboratory (NOAA/GML; https://gml.noaa.gov/ccgg/trends/; last access February 2025).

The seasonal peak in GPP was identified for each pixel based on the monthly distribution through the year following Kelley et al. (2013), whereby each month is represented by a vector in the complex plane whose direction (θ_m_) corresponds to the time of year, and length corresponds to the magnitude for that month:

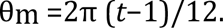

where θm is the direction corresponding to month *t*, with month 1 arbitrarily set to an angle of zero. Peak GPP was considered as month 1 and the seasonal climatologies of both VPD and GPP were aligned to this starting point.

The new set of layers was classified using the X-means clustering algorithm (Fraley & Raftery, 1998; Pelleg & Moore, 2000). X-means is a variant of k-means clustering that refines clustering by repeated sub-division until an optimal number of clusters is obtained, thus removing the need to specify the number of clusters *a priori*. We set the maximum number of clusters to 100, and the optimal number of clusters was then determined automatically using the Bayesian Information Criterion (BIC) recursively. We made two runs, at 10km and 50 km resolution, to investigate whether differences in pixel size influenced the clusters identified.

We examined the characteristics of the clusters in terms of their fire properties, using seven metrics: average fire size (km^-2^), burnt area fraction (%), number of ignitions (km^-2^), average fire speed (km/day^-1^), average fire duration (days), carbon emission (gCm^-2^) and fire season concentration (unitless). Monthly data on average fire size, number of ignitions, average fire speed and average fire duration at 0.25° resolution were obtained from the updated version of the Fire Atlas (Andela et al., 2019), which provides information on fire properties up to mid-2023 (ESA World Fire Atlas, https://s3wfa.esa.int/; Andela and Jones, 2024). Burnt area data were obtained from the MODIS MCD64A1 dataset (Giglio et al., 2006) at a resolution of 0.5°. Fire seasonal concentration was derived from the burnt area data following the same method as the seasonal peak in GPP, described above. Thus, if wildfire is spread over all months then the seasonal concentration is equal to zero whereas it will be equal to 1 if wildfire is concentrated in a single month. Carbon emissions data were obtained from GFED5 (Chen et al., 2023) with an original resolution of 0.25°.

For consistency with the VPD and GPP data, we used fire data covering the interval 2003 to 2020. The datasets were aggregated to a common resolution of 0.5° using bilinear interpolation. The monthly data were averaged over the study period (2003–2020). The cluster data were also aggregated to 0.5° using the modal cluster type. Fire property layers were then stacked with the cluster data to extract fire property information for each grid cell.

We used Hotellings *T^2^* statistic (Hotelling, 1931; Wilks, 2011) to assess the overall dissimilarity between the clusters across all the 7 fire properties: average fire size, burnt area fraction, number of ignitions, average fire speed, average fire duration, carbon emission and fire season concentration. Pairwise Wilcoxon tests (Wilcoxon 1992; Conover 1999) were then applied to estimate the significance of differences between each pair of clusters for each fire property. Additionally, we identified significant differences between a specific cluster and all other clusters for a given fire property to determine which properties best distinguish that cluster from the rest.

We used generalized linear models (GLMs) to examine the degree to which variables relating to vegetation type and human activity influenced within-cluster variation in fire properties. For each cluster, a GLM was fitted using a quasi-Poisson link function for each fire property against annual GPP (gCyr^-1^), maximum VPD (Pa), fractional grass, shrub and tree cover, fractional cropland and pasture cover, road density and population density. We used simulated GPP from the P Model, maximum VPD from ERA5-Land (Muñoz-Sabater, 2019), fractional grass, shrub and tree cover from ESA CCI landcover (Li et al 2018), fractional cropland and pasture cover and population density from HYDE 3.3 (Klein Goldewijk, 2023), and road density from the GRIP dataset (Meijer et al., 2018). The vegetation indices (tree cover, shrub cover, grass cover) were obtained by converting the ESA CCI landcover types to fractional cover following Poulter *et al*. (2015) based on the conversion table from Forkel *et al*. (2017). Cropland area and pasture area were converted to fractional data by dividing by the area of the grid cell to remove the bias caused by differences in grid cell size. We calculated an average value for the interval 2003-2020 for population density, cropland area, and pasture area; the GRIP dataset provides a single snapshot for 2018. GPP and VPD were log transformed for these models, but no transformation was applied to the other variables. The analyses were performed at 0.5° resolution. We aggregated all the data from their original resolution to 0.5° using bilinear interpolation. We also performed a second GLM analysis to assess the influence of environmental variability on the vegetation and human predictors by excluding GPP and VPD. We ranked the importance of the variables in each GLM model at driving intra-cluster variability using *t-values.* Given that the sample size differs between clusters, the *t-values* were standardized but preserving the sign to reflect whether the variable had a positive or negative effect on a given fire property. Thus, the standardised values vary between -1 and 1. Given the large sample sizes (Table 1), the *t-values* were set to zero when the fitted relationship was not significant (p<0.001). The chi-squared *p-value* of each GLM (p<0.01) was calculated using the *broom* package in R, and R^2^ values of significant models are reported in bold.

## Data Availability

Climate data from ERA5-Land are available from Copernicus Climate Change Service (C3S) Climate Data Store (CDS). https://doi.org/10.24381/cds.68d2bb30. Soil data from Soilgrids 2.0 are available from https://doi.org/10.5194/soil-7-217-2021. Annual CO_2_ is available from the NOAA Global Monitoring Laboratory at https://gml.noaa.gov/ccgg/trends/. The fAPAR data from Jeong et al. (2024) are available from the authors on request. Fire properties were obtained from The Global Fire Atlas available at https://doi.org/10.5281/zenodo.11400062. The MODIS MCD64A1 burnt area data set is available from https://lpdaac.usgs.gov/products/mcd64a1v061/. Population density, cropland area, and pasture area were obtained from the HYDE 3.3 database, available at http://doi.org/10.24416/UU01-67UHB4. Road density data were obtained from the Global Roads Inventory Project (GRIP) dataset available at http://www.globio.info/download-grip-dataset. The global data set of simulated GPP and the aggregated clusters are available at https://figshare.com/s/3904aa4a3ca5111e5856.

## Code Availability

The P model code is available from https://doi.org/10.5281/zenodo.8366848 The SPLASH code is available from https://github.com/dsval/rsplash. The X-means clustering code is available from https://code.earthengine.google.com/ac5b1cf3e4cadb5507d874da5c679215. The chi-squared *p-value* of each GLM (p<0.01) was calculated using the *broom* package in R.

## Acknowledgements

We thank Matthias Forkel, Matt Forrest and Sander Veraverbeke for feedback on an earlier version of this paper, and our colleagues at the Leverhulme Centre for Wildfires Environment and Society for discussions. This research received financial support through Schmidt Sciences, LLC (SPH, OH, DSC, ICP) and the Leverhulme Centre for Wildfires Environment and Society (YS, DS).

## Acknowledgements

This work is a contribution to the LEMONTREE (Land Ecosystem Models based On New Theory, obseRvations and ExperimEnts) project, and to the Leverhulme Centre for Wildfires, Environment and Society. This research received support through Schmidt Sciences, LLC (Grant number 355: ICP, SPH, OH, DSC, YS). DS acknowledges support from the Leverhulme Centre for Wildfires, Environment and Society (Grant number: RC-2018-023).

## Author Contributions

ICP and SPH developed the underlying theory. ICP, SPH and OH designed the study. OH, DSC, YS and DS carried out data analyses and produced the figures. SPH wrote the first draft of the text and all authors contributed to the final draft.

