## Supplementary Material for "An eco-evolutionary approach to defining wildfire regimes"

**An eco-evolutionary approach to defining wildfire regimes: Supplementary Material**

This Supplementary contains the following Figures and Tables:

Supplementary Figure 1: Global map showing the region occupied by Cluster 1, as derived from X-means clustering based on in the magnitude and seasonal phasing of gross primary production (GPP) and vapour pressure deficit (VPD) of data at 10 km resolution.

Supplementary Figure 2: Global map showing the region occupied by Cluster 2, as derived from X-means clustering based on in the magnitude and seasonal phasing of gross primary production (GPP) and vapour pressure deficit (VPD) of data at 10 km resolution.

Supplementary Figure 3: Global map showing the region occupied by Cluster 3, as derived from X-means clustering based on in the magnitude and seasonal phasing of gross primary production (GPP) and vapour pressure deficit (VPD) of data at 10 km resolution.

Supplementary Figure 4: Global map showing the region occupied by Cluster 4, as derived from X-means clustering based on in the magnitude and seasonal phasing of gross primary production (GPP) and vapour pressure deficit (VPD) of data at 10 km resolution.

Supplementary Figure 5: Global map showing the region occupied by Cluster 5, as derived from X-means clustering based on in the magnitude and seasonal phasing of gross primary production (GPP) and vapour pressure deficit (VPD) of data at 10 km resolution.

Supplementary Figure 6: Global map showing the region occupied by Cluster 6, as derived from X-means clustering based on in the magnitude and seasonal phasing of gross primary production (GPP) and vapour pressure deficit (VPD) of data at 10 km resolution.

Supplementary Figure 7: Global map showing the region occupied by Cluster 7, as derived from X-means clustering based on in the magnitude and seasonal phasing of gross primary production (GPP) and vapour pressure deficit (VPD) of data at 10 km resolution.

Supplementary Figure 8: Global map showing the region occupied by Cluster 8, as derived from X-means clustering based on in the magnitude and seasonal phasing of gross primary production (GPP) and vapour pressure deficit (VPD) of data at 10 km resolution.

Supplementary Figure 9: Global map showing the region occupied by Cluster 9, as derived from X-means clustering based on in the magnitude and seasonal phasing of gross primary production (GPP) and vapour pressure deficit (VPD) of data at 10 km resolution.

Supplementary Figure 10: Global map showing the region occupied by Cluster 10, as derived from X-means clustering based on in the magnitude and seasonal phasing of gross primary production (GPP) and vapour pressure deficit (VPD) of data at 10 km resolution.

Supplementary Figure 11: Global map showing the region occupied by Cluster 11, as derived from X-means clustering based on in the magnitude and seasonal phasing of gross primary production (GPP) and vapour pressure deficit (VPD) of data at 10 km resolution.

Supplementary Figure 12: Global map showing the region occupied by Cluster 12, as derived from X-means clustering based on in the magnitude and seasonal phasing of gross primary production (GPP) and vapour pressure deficit (VPD) of data at 10 km resolution.

Supplementary Figure 13: Global map showing the region occupied by Cluster 13, as derived from X-means clustering based on in the magnitude and seasonal phasing of gross primary production (GPP) and vapour pressure deficit (VPD) of data at 10 km resolution.

Supplementary Figure 14: Global map showing the region occupied by Cluster 14, as derived from X-means clustering based on in the magnitude and seasonal phasing of gross primary production (GPP) and vapour pressure deficit (VPD) of data at 10 km resolution.

Supplementary Figure 15: Global map showing the region occupied by Cluster 15, as derived from X-means clustering based on in the magnitude and seasonal phasing of gross primary production (GPP) and vapour pressure deficit (VPD) of data at 10 km resolution.

Supplementary Figure 16: Global map showing the region occupied by Cluster 16, as derived from X-means clustering based on in the magnitude and seasonal phasing of gross primary production (GPP) and vapour pressure deficit (VPD) of data at 10 km resolution.

Supplementary Figure 17: Global map showing the region occupied by Cluster 17, as derived from X-means clustering based on in the magnitude and seasonal phasing of gross primary production (GPP) and vapour pressure deficit (VPD) of data at 10 km resolution.

Supplementary Figure 18: Global map showing the region occupied by Cluster 18, as derived from X-means clustering based on in the magnitude and seasonal phasing of gross primary production (GPP) and vapour pressure deficit (VPD) of data at 10 km resolution.

Supplementary Figure 19: Regions of similar patterns in the magnitude and seasonal phasing of gross primary production (GPP) and vapour pressure deficit (VPD) based on X-means clustering of data at 50 km resolution.

Supplementary Figure 20: Cross-correlations between wildfire properties: average burnt area ($km^{-2}$), average fire size ($km^{-2}$), average number of ignitions ($km^{-2}$), average fire speed ($km/day^{-1}$), average fire duration (days), average seasonal concentration (unitless), and average carbon emission ($gC/m^{-2}$).

Supplementary Table 1: Significant differences between a specific cluster and all other clusters for a given fire property, where ✓ indicates significant difference $p < 0.01$.

Supplementary Table 2: Summary of results for the generalised linear model (GLM) of the within-cluster variability in fire properties as a function of fractional grass cover (G), fractional shrub cover (S), fractional tree cover (T), fractional cropland cover (C), fractional pasture cover (P), road density (R) and population density (Pn). $R^2$ values for significant models are given in **bold**; the significant factors (F) are given in order of importance and colour-coded to indicate whether their impact on a fire property is positive (in red) or negative (in blue).

Supplementary Table 3: Results for the generalised linear model (GLM) of the within-cluster variability in burnt area as a function of the annual gross primary production (GPP, $gCyr^{-1}$), maximum vapour pressure deficit (VPD, Pa), fractional grass cover (grass), fractional shrub cover (shrub), fractional tree cover tree), fractional cropland cover (crop), fractional pasture cover (pasture), road density (roads) and population density (popn). $R^2$ values for significant models are given in **bold.** Given that the sample size differs between clusters, the *t-values* were standardized but preserving the sign to reflect whether the variable had a positive or negative effect on burnt area. Thus, the standardised values vary between -1 and 1. Only significant *t-values* are shown.

Supplementary Table 4: Results for the generalised linear model (GLM) of the within-cluster variability in number of ignitions as a function of the annual gross primary production (GPP, $gCyr^{-1}$), maximum VPD (VPD, Pa), fractional grass cover (grass), fractional shrub cover (shrub), fractional tree cover tree), fractional cropland cover (crop), fractional pasture cover (pasture), road density (roads) and population density (popn). $R^2$ values for significant models are given in **bold.** Given that the sample size differs between clusters, the *t-values* were standardized but preserving the sign to reflect whether the variable had a positive or negative effect on number of ignitions. Thus, the standardised values vary between -1 and 1. Only significant *t-values* are shown.

Supplementary Table 5: Results for the generalised linear model (GLM) of the within-cluster variability in average fire size as a function of the annual gross primary production (GPP, $gCyr^{-1}$), maximum VPD (VPD, Pa), fractional grass cover (grass), fractional shrub cover (shrub), fractional tree cover tree), fractional cropland cover (crop), fractional pasture cover (pasture), road density (roads) and population density (popn). $R^2$ values for significant models are given in **bold.** Given that the sample size differs between clusters, the *t-values* were standardized but preserving the sign to reflect whether the variable had a positive or negative effect on number of ignitions. Thus, the standardised values vary between -1 and 1. Only significant *t-values* are shown.

Supplementary Table 6: Results for the generalised linear model (GLM) of the within-cluster variability in average fire speed as a function of the annual gross primary production (GPP, $gCyr^{-1}$), maximum VPD (VPD, Pa), fractional grass cover (grass), fractional shrub cover (shrub), fractional tree cover tree), fractional cropland cover (crop), fractional pasture cover (pasture), road density (roads) and population density (popn). $R^2$ values for significant models are given in **bold.** Given that the sample size differs between clusters, the *t-values* were standardized but preserving the sign to reflect whether the variable had a positive or negative effect on number of ignitions. Thus, the standardised values vary between -1 and 1. Only significant *t-values* are shown.

Supplementary Table 7: Results for the generalised linear model (GLM) of the within-cluster variability in average fire duration as a function of the annual gross primary production (GPP, $gCyr^{-1}$), maximum VPD (VPD, Pa), fractional grass cover (grass), fractional shrub cover (shrub), fractional tree cover tree), fractional cropland cover (crop), fractional pasture cover (pasture), road density (roads) and population density (popn). $R^2$ values for significant models are given in **bold.** Given that the sample size differs between clusters, the *t-values* were standardized but preserving the sign to reflect whether the variable had a positive or negative effect on number of ignitions. Thus, the standardised values vary between -1 and 1. Only significant *t-values* are shown.

Supplementary Table 8: Results for the generalised linear model (GLM) of the within-cluster variability in carbon emissions as a function of the annual gross primary production (GPP, $gCyr^{-1}$), maximum VPD (VPD, Pa), fractional grass cover (grass), fractional shrub cover (shrub), fractional tree cover tree), fractional cropland cover (crop), fractional pasture cover (pasture), road density (roads) and population density (popn). $R^2$ values for significant models are given in **bold.** Given that the sample size differs between clusters, the *t-values* were standardized but preserving the sign to reflect whether the variable had a positive or negative effect on number of ignitions. Thus, the standardised values vary between -1 and 1. Only significant *t-values* are shown.

Supplementary Table 9: Results for the generalised linear model (GLM) of the within-cluster variability in fire seasonal concentration as a function of the annual gross primary production (GPP, $gCyr^{-1}$), maximum VPD (VPD, Pa), fractional grass cover (grass), fractional shrub cover (shrub), fractional tree cover tree), fractional cropland cover (crop), fractional pasture cover (pasture), road density (roads) and population density (popn). $R^2$ values for significant models are given in **bold.** Given that the sample size differs between clusters, the *t-values* were standardized but preserving the sign to reflect whether the variable had a positive or negative effect on number of ignitions. Thus, the standardised values vary between -1 and 1. Only significant *t-values* are shown.

Supplementary Figure 1: Global map showing the region occupied by Cluster 1, as derived from X-means clustering based on in the magnitude and seasonal phasing of gross primary production (GPP) and vapour pressure deficit (VPD) of data at 10 km resolution.

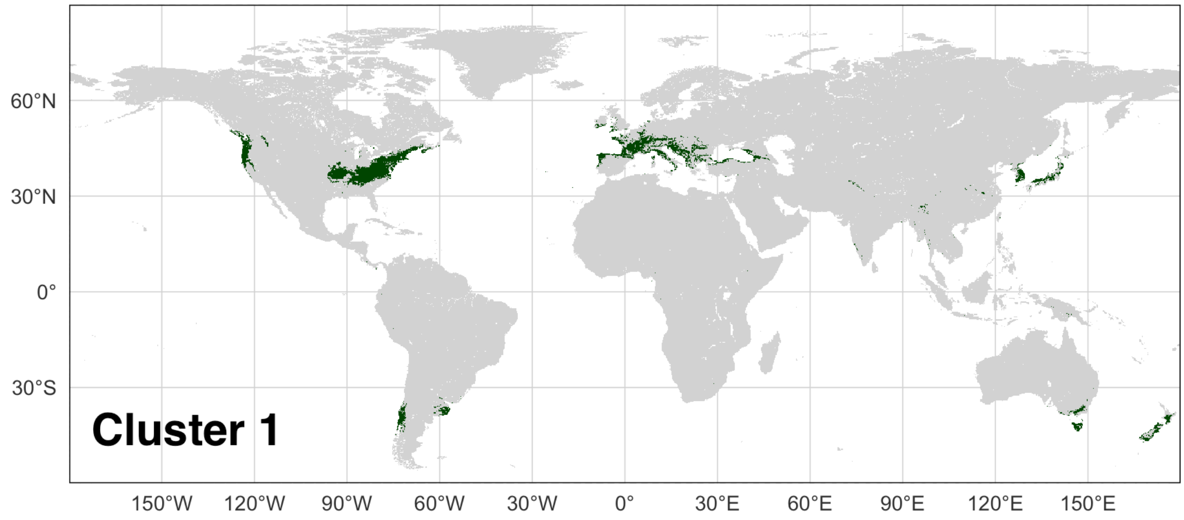


Supplementary Figure 2: Global map showing the region occupied by Cluster 2, as derived from X-means clustering based on in the magnitude and seasonal phasing of gross primary production (GPP) and vapour pressure deficit (VPD) of data at 10 km resolution.

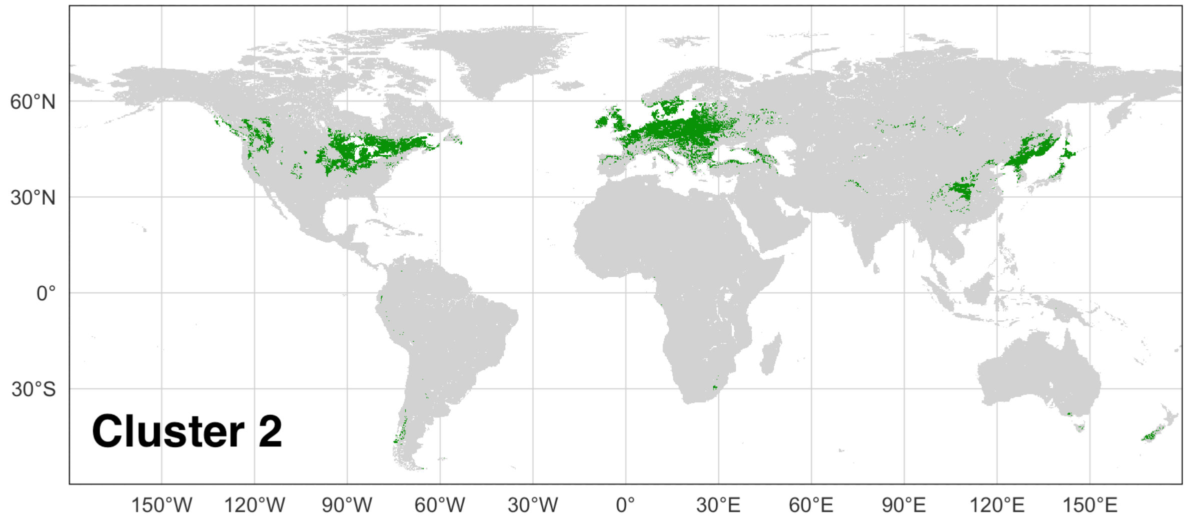

Supplementary Figure 3: Global map showing the region occupied by Cluster 3, as derived from X-means clustering based on in the magnitude and seasonal phasing of gross primary production (GPP) and vapour pressure deficit (VPD) of data at 10 km resolution.

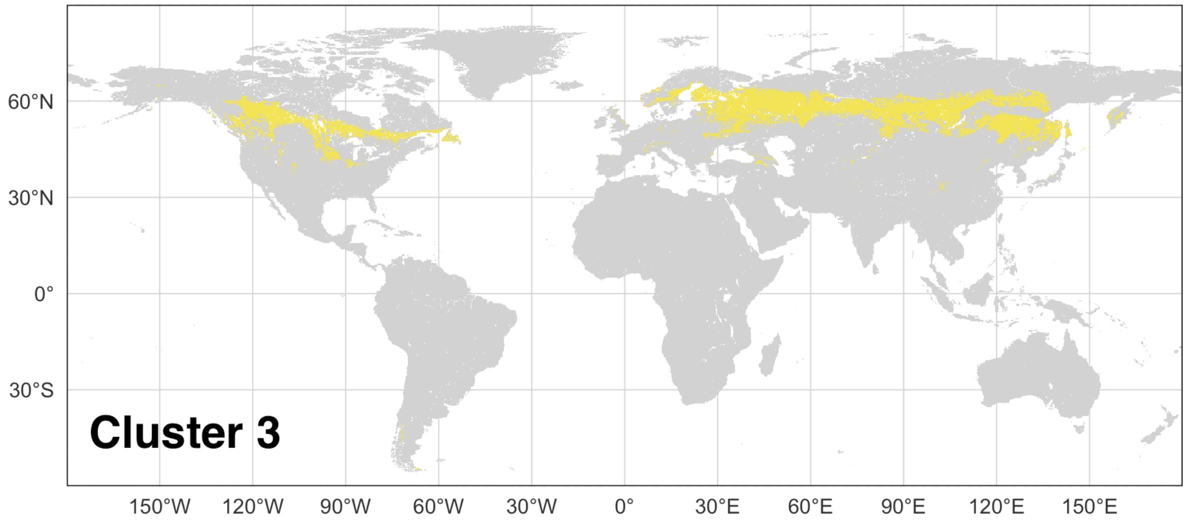


Supplementary Figure 4: Global map showing the region occupied by Cluster 4, as derived from X-means clustering based on in the magnitude and seasonal phasing of gross primary production (GPP) and vapour pressure deficit (VPD) of data at 10 km resolution.

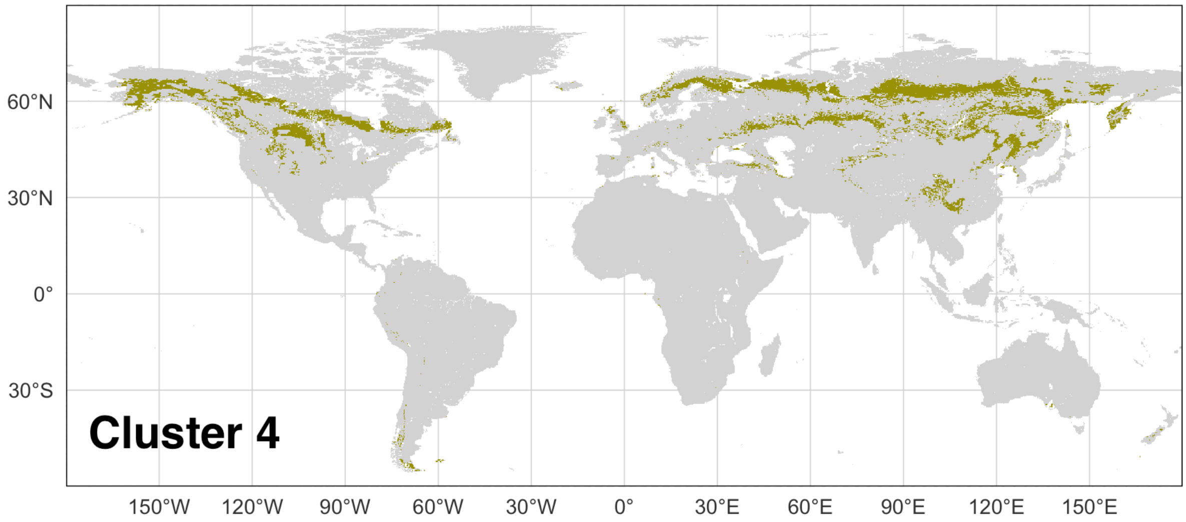

Supplementary Figure 5: Global map showing the region occupied by Cluster 5, as derived from X-means clustering based on in the magnitude and seasonal phasing of gross primary production (GPP) and vapour pressure deficit (VPD) of data at 10 km resolution.

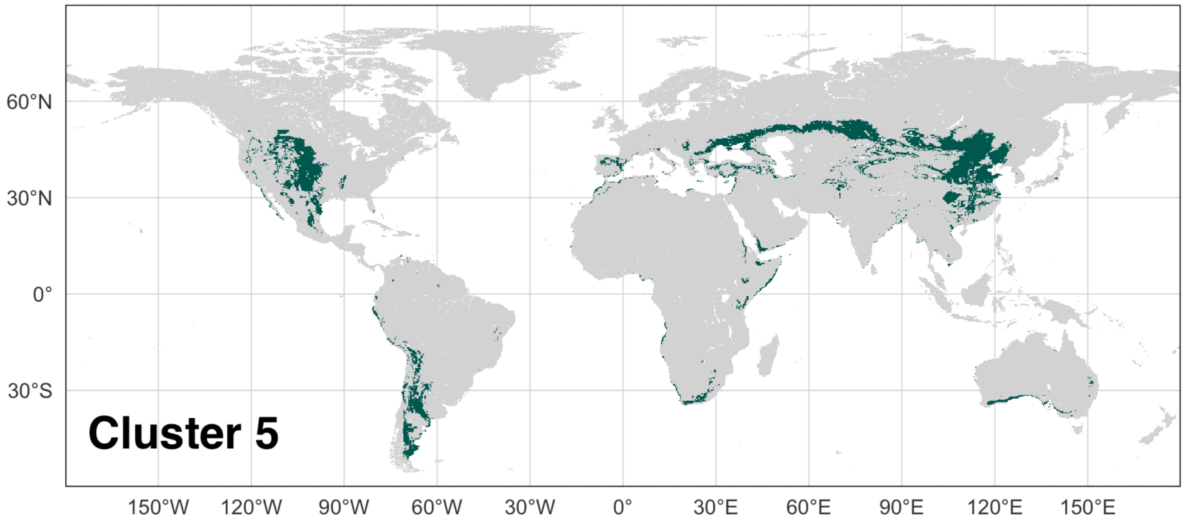


Supplementary Figure 6: Global map showing the region occupied by Cluster 6, as derived from X-means clustering based on in the magnitude and seasonal phasing of gross primary production (GPP) and vapour pressure deficit (VPD) of data at 10 km resolution.

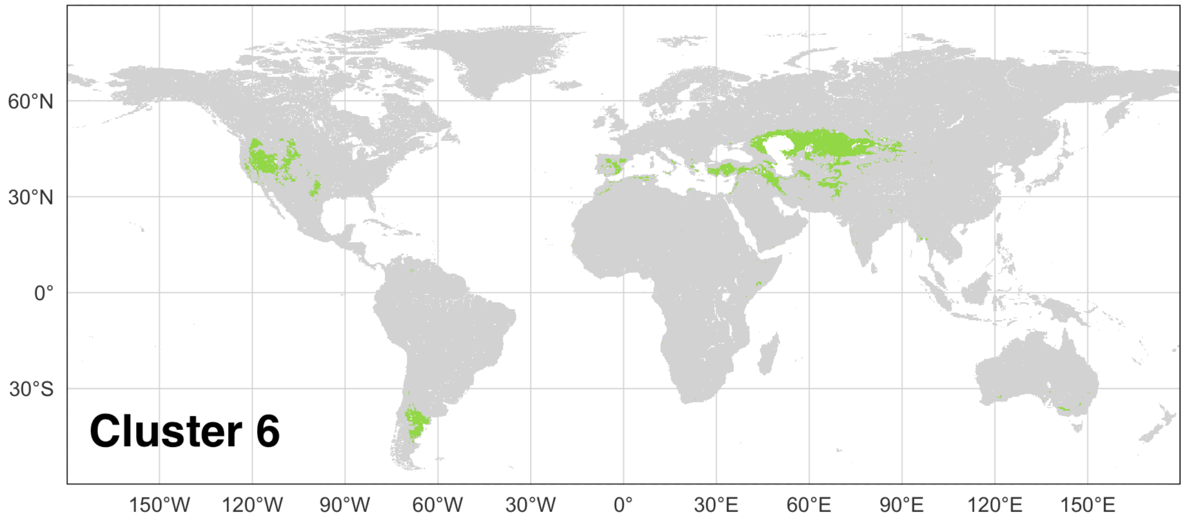

Supplementary Figure 7: Global map showing the region occupied by Cluster 7, as derived from X-means clustering based on in the magnitude and seasonal phasing of gross primary production (GPP) and vapour pressure deficit (VPD) of data at 10 km resolution.

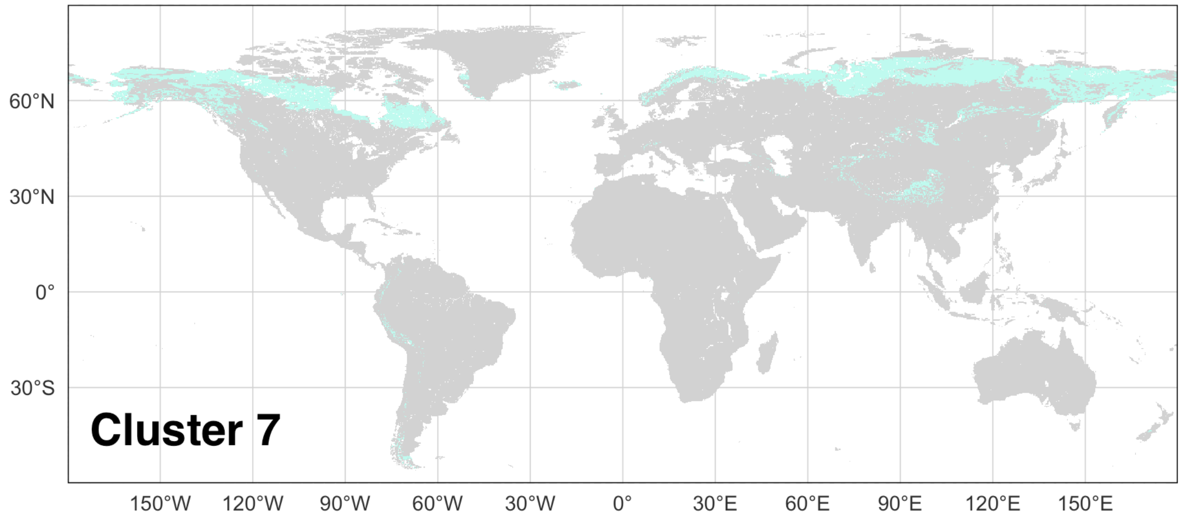


Supplementary Figure 8: Global map showing the region occupied by Cluster 8, as derived from X-means clustering based on in the magnitude and seasonal phasing of gross primary production (GPP) and vapour pressure deficit (VPD) of data at 10 km resolution.

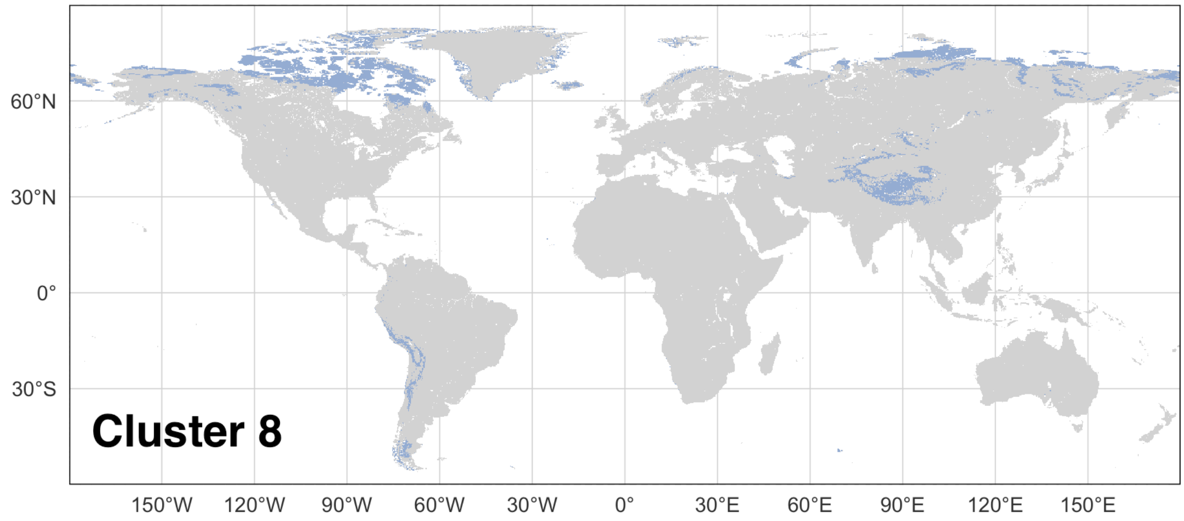

Supplementary Figure 9: Global map showing the region occupied by Cluster 9, as derived from X-means clustering based on in the magnitude and seasonal phasing of gross primary production (GPP) and vapour pressure deficit (VPD) of data at 10 km resolution.

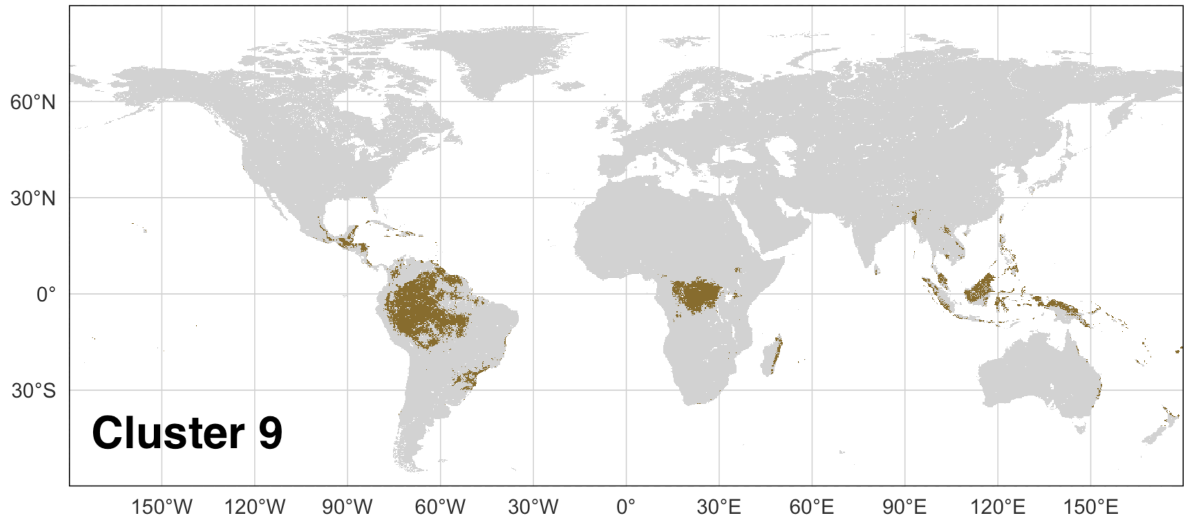


Supplementary Figure 10: Global map showing the region occupied by Cluster 10, as derived from X-means clustering based on in the magnitude and seasonal phasing of gross primary production (GPP) and vapour pressure deficit (VPD) of data at 10 km resolution.

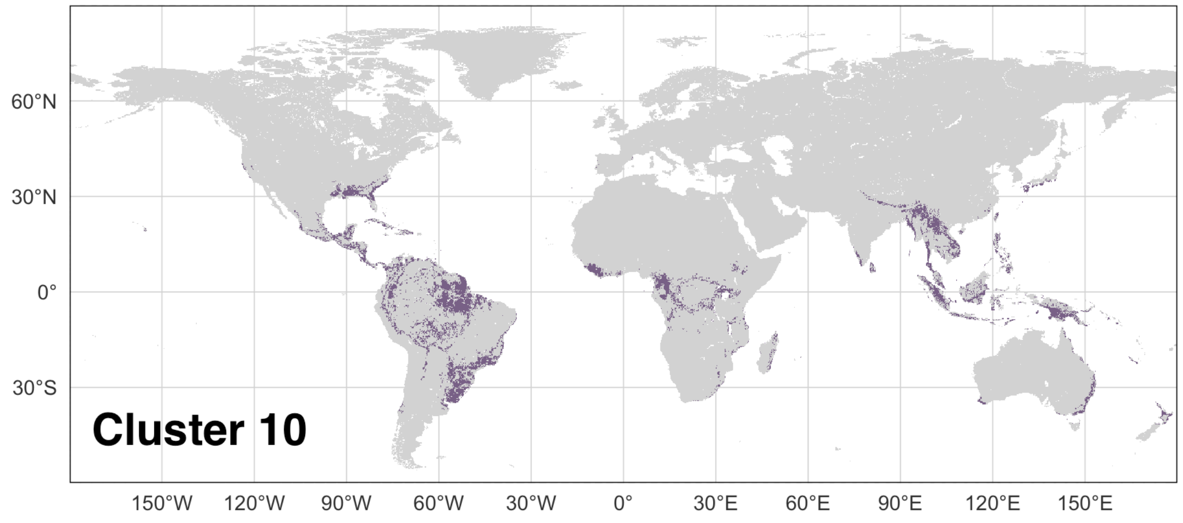

Supplementary Figure 11: Global map showing the region occupied by Cluster 11, as derived from X-means clustering based on in the magnitude and seasonal phasing of gross primary production (GPP) and vapour pressure deficit (VPD) of data at 10 km resolution.

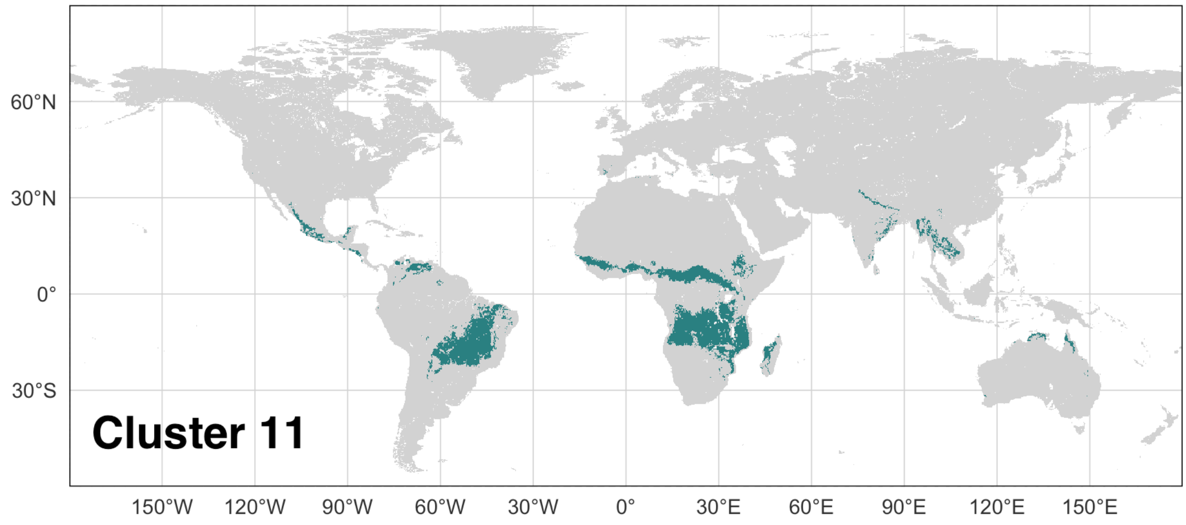


Supplementary Figure 12: Global map showing the region occupied by Cluster 12, as derived from X-means clustering based on in the magnitude and seasonal phasing of gross primary production (GPP) and vapour pressure deficit (VPD) of data at 10 km resolution.

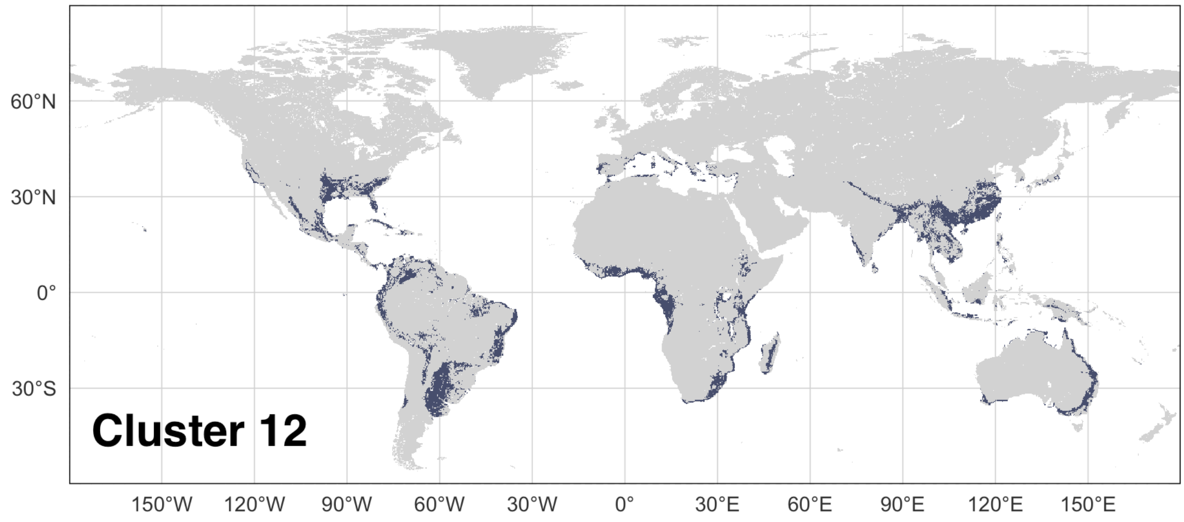

Supplementary Figure 13: Global map showing the region occupied by Cluster 13, as derived from X-means clustering based on in the magnitude and seasonal phasing of gross primary production (GPP) and vapour pressure deficit (VPD) of data at 10 km resolution.

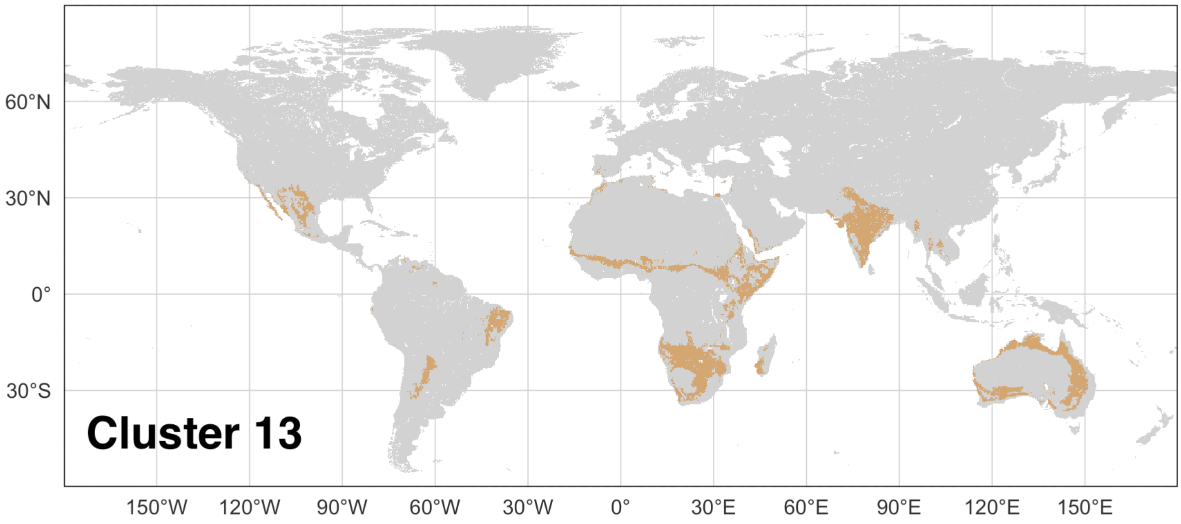


Supplementary Figure 14: Global map showing the region occupied by Cluster 14, as derived from X-means clustering based on in the magnitude and seasonal phasing of gross primary production (GPP) and vapour pressure deficit (VPD) of data at 10 km resolution.

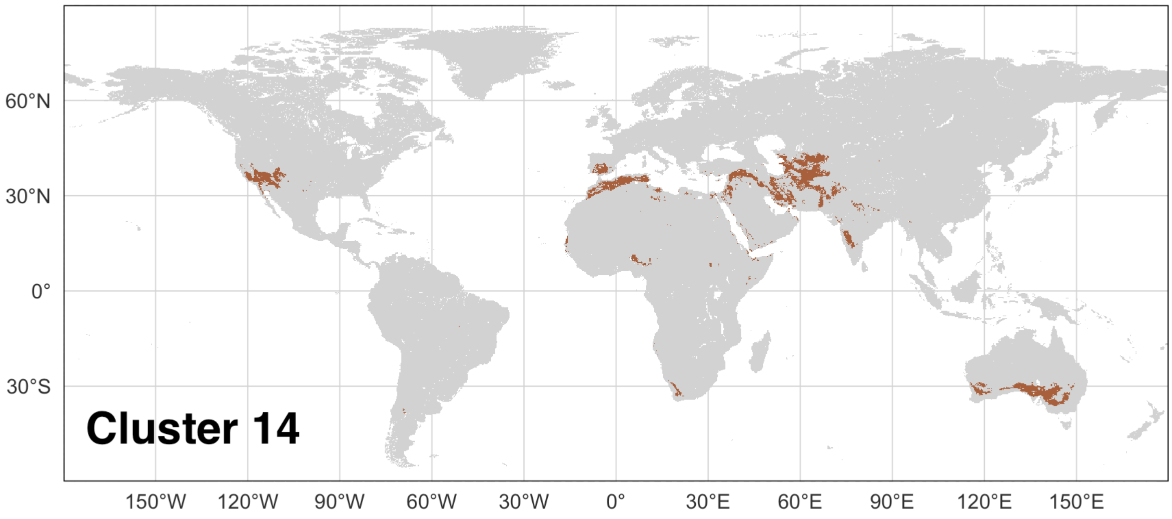

Supplementary Figure 15: Global map showing the region occupied by Cluster 15, as derived from X-means clustering based on in the magnitude and seasonal phasing of gross primary production (GPP) and vapour pressure deficit (VPD) of data at 10 km resolution.

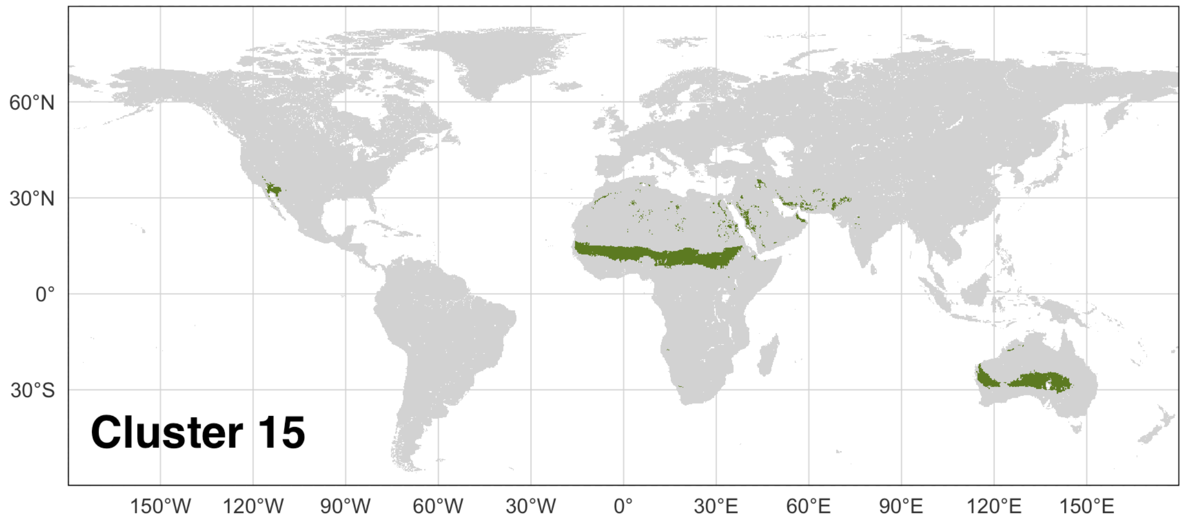


Supplementary Figure 16: Global map showing the region occupied by Cluster 16, as derived from X-means clustering based on in the magnitude and seasonal phasing of gross primary production (GPP) and vapour pressure deficit (VPD) of data at 10 km resolution.

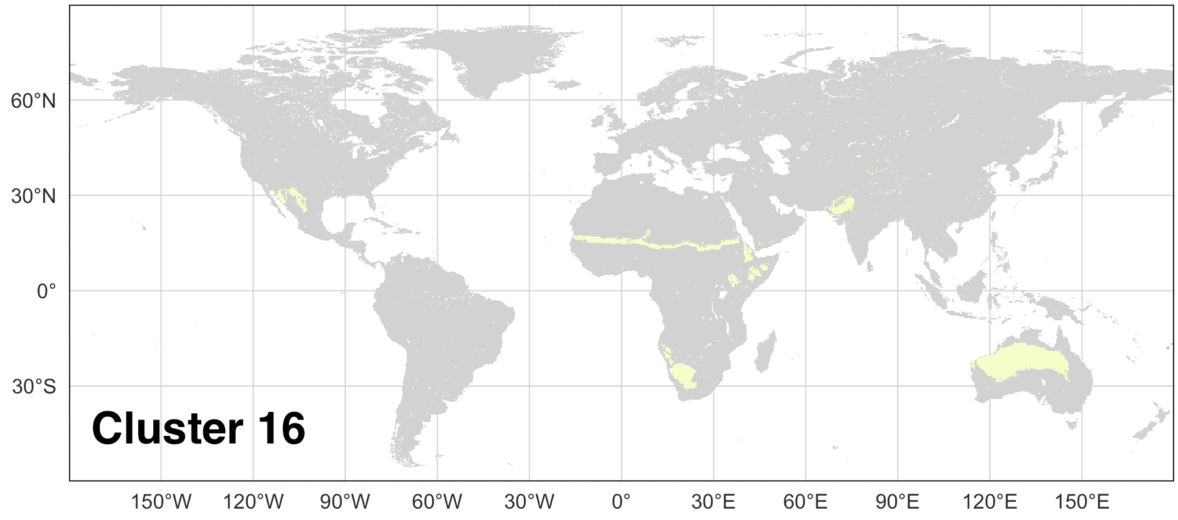

Supplementary Figure 17: Global map showing the region occupied by Cluster 17, as derived from X-means clustering based on in the magnitude and seasonal phasing of gross primary production (GPP) and vapour pressure deficit (VPD) of data at 10 km resolution.

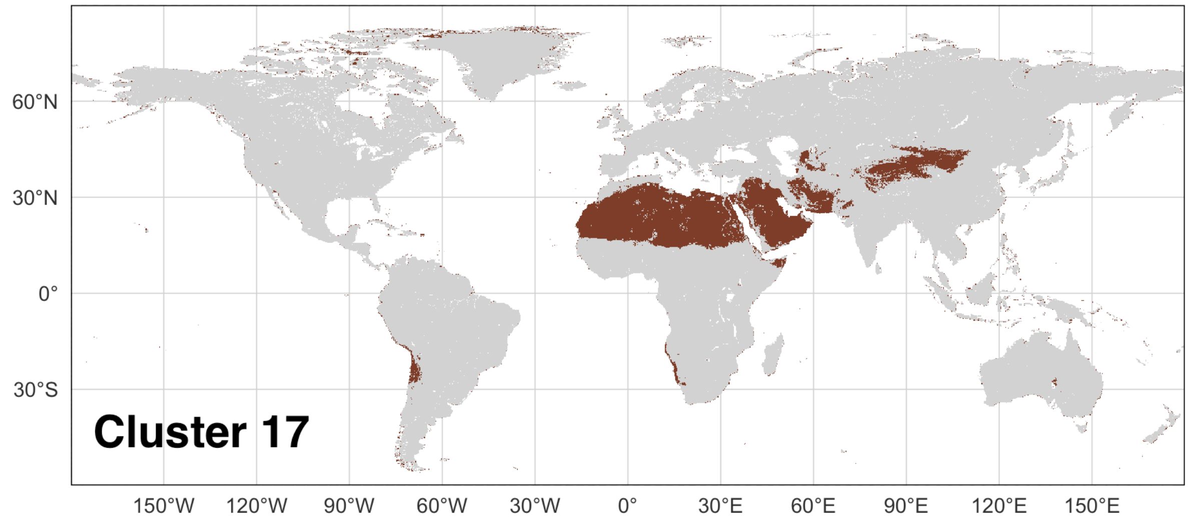


Supplementary Figure 18: Global map showing the region occupied by Cluster 18, as derived from X-means clustering based on in the magnitude and seasonal phasing of gross primary production (GPP) and vapour pressure deficit (VPD) of data at 10 km resolution.

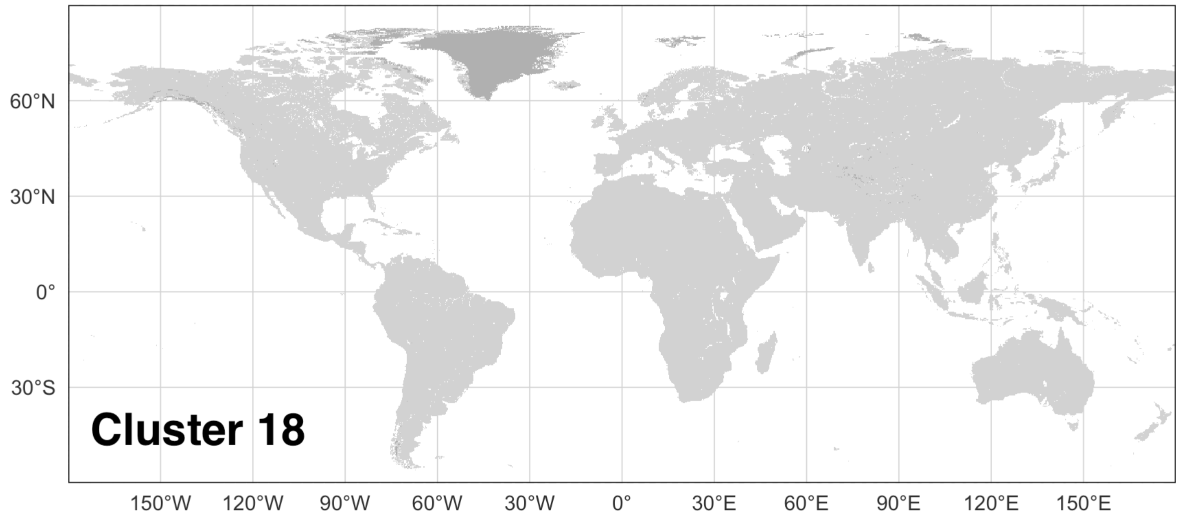

Supplementary Figure 19: Regions of similar patterns in the magnitude and seasonal phasing of gross primary production (GPP) and vapour pressure deficit (VPD) based on X-means clustering of data at 50 km resolution.

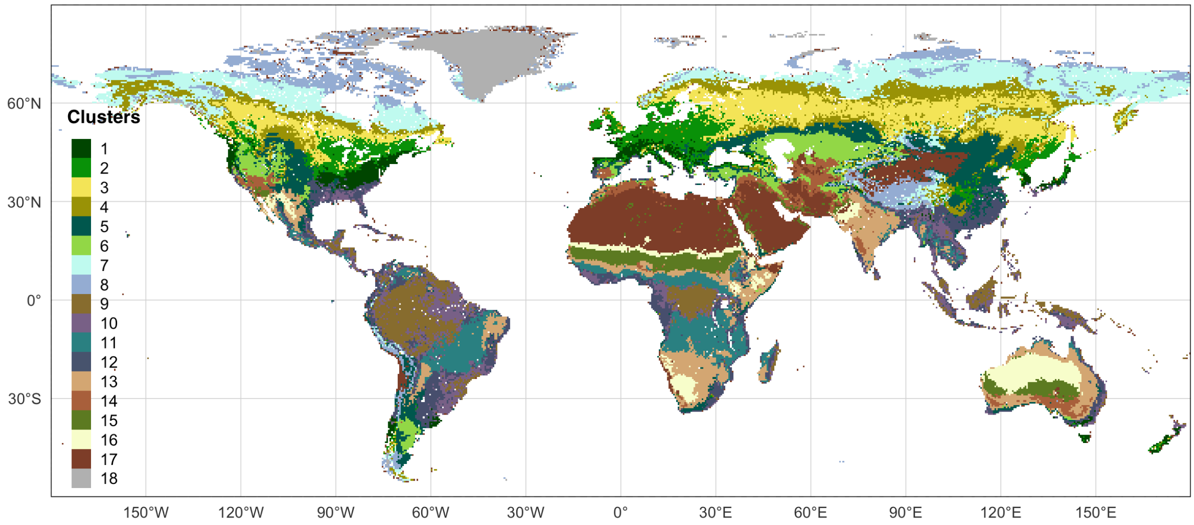


Supplementary Figure 20. Cross-correlations between wildfire properties: average burnt area ($km^{-2}$), average fire size ($km^{-2}$), average number of ignitions ($km^{-2}$), average fire speed ($km/day^{-1}$), average fire duration (days), average seasonal concentration (unitless), and average carbon emission ($gC/m^{-2}$).

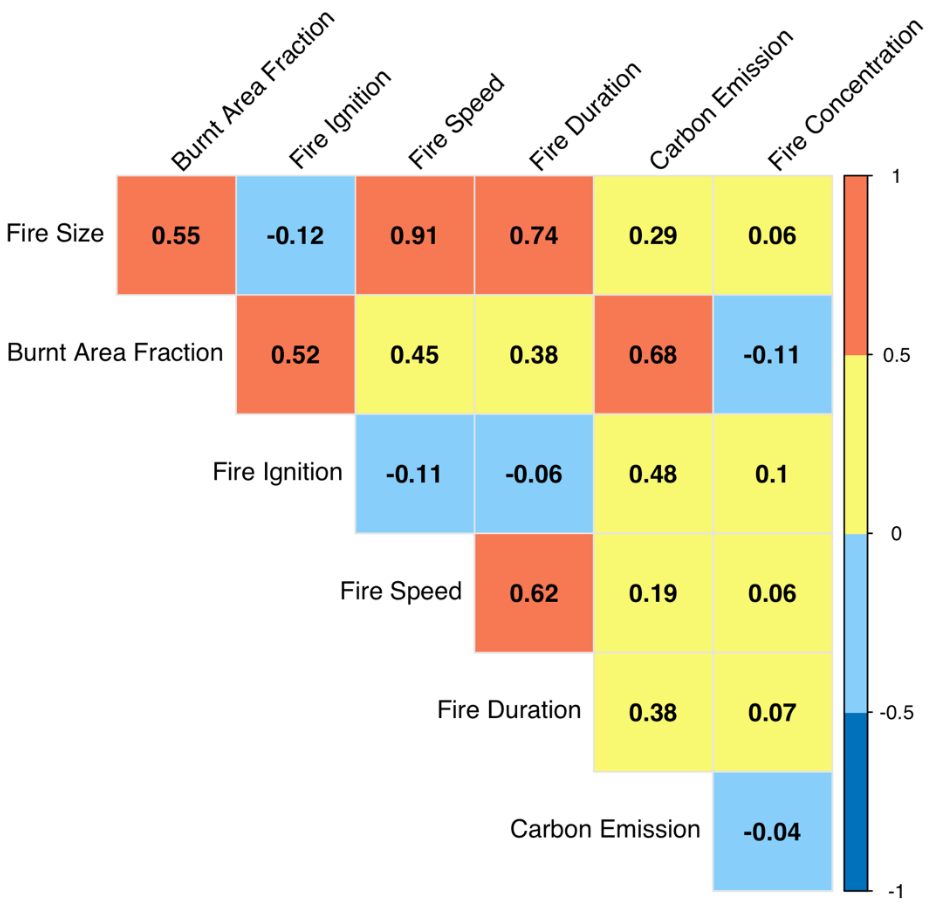
Burnt Area Fraction
Fire Ignition
Fire Speed
Fire Duration
Carbon Emission
Fire Concentration
Fire Size
0.55
-0.12
0.91
0.74
0.29
0.06
Burnt Area Fraction
0.52
0.45
0.38
0.68
-0.11
Fire Ignition
-0.11
-0.06
0.48
0.1
Fire Speed
0.62
0.19
0.06
Fire Duration
0.38
0.07
Carbon Emission
-0.04
1
0.5
0
-0.5
-1

Supplementary Table 1: Significant differences between a specific cluster and all other clusters for a given fire property, where ✓ indicates significant difference $p < 0.01$.

| Cluster | Fire size | Burnt area fraction | Fire ignition | Fire speed | Fire duration | Carbon emission | Fire concentration |
|---|---|---|---|---|---|---|---|
| 1 | | | | | | | |
| 2 | | | | | | ✓ | |
| 3 | | ✓ | | | | | |
| 4 | ✓ | ✓ | | | | | ✓ |
| 5 | | ✓ | | | | | |
| 6 | | | | ✓ | | | |
| 7 | | | | | | | |
| 8 | | | | | | ✓ | |
| 9 | ✓ | | | | ✓ | | |
| 10 | ✓ | | | | | ✓ | |
| 11 | | ✓ | ✓ | ✓ | | ✓ | |
| 12 | | ✓ | | | | | |
| 13 | ✓ | | | | | | ✓ |
| 14 | | ✓ | ✓ | | | ✓ | |
| 15 | | ✓ | | ✓ | | | |
| 16 | ✓ | | ✓ | ✓ | ✓ | | ✓ |
| 17 | | | ✓ | | | | |
| 18 | | | | | | | |

Supplementary Table 2: Summary of results for the generalised linear model (GLM) of the within-cluster variability in fire properties as a function of fractional grass cover (G), fractional shrub cover (S), fractional tree cover (T), fractional cropland cover (C), fractional pasture cover (P), road density (R) and population density (Pn). $R^2$ values for significant models are given in **bold**; the significant factors (F) are given in order of importance and colour-coded to indicate whether their impact on a fire property is positive (in red) or negative (in blue).

| | BA | | Ignitions | | Size | | Speed | | Duration | | Emissions | | Concentration | |
|---|---|---|---|---|---|---|---|---|---|---|---|---|---|---|
| cluster | R2 | *F* | R2 | *F* | R2 | *F* | R2 | *F* | R2 | *F* | R2 | *F* | R2 | *F* |
| 1 | 0.2 | *ns* | **0.03** | *ns* | **0.22** | *ns* | **0.06** | *ns* | **0.07** | T | **0.42** | S G T P | **0.06** | C P R |
| 2 | 0.21 | *ns* | **0.06** | R S | **0.21** | S | **0.13** | S | **0.03** | S | 0.12 | *ns* | **0.04** | R C |
| 3 | **0.14** | G T R C S | **0.08** | G P T | 0.13 | *ns* | **0.09** | R G | **0.15** | S R | 0.1 | *ns* | **0.2** | C T R S |
| 4 | **0.14** | C P R S Pn | **0.07** | S P C G R | 0.14 | *ns* | **0.11** | S C | **0.22** | S T P | **0.11** | S C T R | **0.36** | C T R Pn |
| 5 | **0.1** | C G P | **0.34** | C G S | 0.32 | *ns* | **0.21** | C P T | **0.05** | T S | **0.16** | C S T | **0.04** | C |
| 6 | 0.27 | *ns* | **0.23** | C S R G | 0.33 | *ns* | **0.32** | C G R T | **0.13** | C G S T | **0.14** | C G S | **0.02** | C |
| 7 | **0.18** | T S G | **0.04** | S P C | 0.14 | *ns* | **0.11** | T G S R | **0.11** | T G S R | **0.17** | T S G | **0.04** | R C |
| 8 | **0.12** | S T | **0.24** | R T | 0.09 | *ns* | 0.02 | | **0.17** | T R S | **0.3** | T S | **0.17** | C R |
| 9 | **0.19** | *ns* | **0.16** | G P S | 0.04 | *ns* | **0.04** | C S T | **0.04** | S P | **0.15** | S T | **0.02** | T S |
| 10 | 0.17 | *ns* | **0.1** | S R T | **0.07** | *ns* | **0.03** | T C | **0.01** | T | **0.15** | S R P | **0.1** | S G T |
| 11 | **0.35** | G T R S C | **0.37** | G T P C R | 0.3 | *ns* | **0.22** | S P R C | **0.18** | G P C S R | **0.44** | T G P R | **0.11** | T G P |
| 12 | 0.26 | *ns* | **0.12** | P R S | **0.17** | S P T C | **0.08** | T S P R C | **0.04** | S P T | **0.09** | S R Pn | **0.05** | T P Pn G S |
| 13 | 0.27 | *ns* | **0.24** | P T C G | 0.48 | *ns* | **0.31** | P T C S | **0.14** | P S C | 0.33 | *ns* | **0.02** | P |
| 14 | 0.35 | P | **0.32** | P C | **0.24** | S | **0.19** | S | **0.09** | S T | **0.46** | P C | **0.02** | R |
| 15 | **0.58** | S T P C G | **0.51** | S T P G C | 0.46 | *ns* | **0.27** | C S P | **0.07** | C | **0.68** | S T P G C | **0.04** | R C Pn |
| 16 | 0.38 | *ns* | **0.09** | C P | 0.41 | *ns* | 0.32 | *ns* | **0.24** | G R T C | 0.26 | *ns* | 0.01 | *ns* |
| 17 | **0.18** | P S C | **0.06** | *ns* | 0.12 | *ns* | **0.11** | G | **0.07** | *ns* | **0.36** | T P C S G | 0.01 | *ns* |
| 18 | **0.59** | S T | 0.4 | *ns* | **0.85** | S | 0.49 | *ns* | 0.42 | *ns* | **0.65** | T | **0.59** | P |

Supplementary Table 3: Results for the generalised linear model (GLM) of the within-cluster variability in burnt area as a function of the annual gross primary production (GPP, $gCyr^{-1}$), maximum vapour pressure deficit (VPD, Pa), fractional grass cover (grass), fractional shrub cover (shrub), fractional tree cover tree), fractional cropland cover (crop), fractional pasture cover (pasture), road density (roads) and population density (popn). $R^2$ values for significant models are given in **bold.** Given that the sample size differs between clusters, the *t-values* were standardized but preserving the sign to reflect whether the variable had a positive or negative effect on burnt area. Thus, the standardised values vary between -1 and 1. Only significant *t-values* are shown.

| **cluster** | **r2** | **t values** | | | | | | | | |
|---|---|---|---|---|---|---|---|---|---|---|
| | | **tree** | **shrub** | **grass** | **popn** | **crop** | **pasture** | **roads** | **GPP** | **VPD** |
| 1 | **0.34** | 0 | 0.480 | 0 | 0 | 0 | 0 | 0 | 0.506 | 1 |
| 2 | **0.30** | 0 | 0 | 0 | 0 | 0 | 0 | 0 | 0 | 0 |
| 3 | **0.24** | -0.394 | 0 | 0.637 | 0 | 0 | 0.188 | -0.162 | 0 | 1 |
| 4 | **0.29** | 0 | 0.263 | -0.206 | 0 | 0 | 0.379 | -0.262 | 0 | 1 |
| 5 | **0.11** | -0.462 | 0 | 0.667 | 0 | 1 | -0.599 | 0 | 0.523 | 0 |
| 6 | **0.31** | 0 | 0 | 0 | 0 | 0 | 0 | 0 | 0 | 0 |
| 7 | **0.31** | 0.547 | 0 | 0 | 0 | 0 | 0 | 0 | 0.288 | 1 |
| 8 | **0.22** | 0 | 0 | 0 | 0 | 0 | 0 | 0 | 1 | 0 |
| 9 | **0.45** | -0.163 | 0 | 0.378 | 0 | 0 | 0 | 0 | -0.617 | 1 |
| 10 | **0.31** | 0 | 0 | 0 | 0 | 0 | 0 | 0 | 0 | 0 |
| 11 | **0.38** | 0.686 | 0.359 | 1 | 0 | 0 | 0.282 | -0.592 | 0 | 0.556 |
| 12 | **0.29** | 0 | 0 | 0 | 0 | 0 | 0 | 0 | 0 | 0 |
| 13 | **0.63** | -0.222 | 0.362 | 0.278 | 0 | 0 | -0.127 | -0.165 | 1 | 0.880 |
| 14 | **0.68** | 0 | 0 | 0 | 0 | 0 | 0 | 0 | 0 | 0 |
| 15 | **0.78** | 0 | 0.321 | 0 | -0.147 | -0.169 | -0.186 | 0 | 1 | 0.218 |
| 16 | 0.68 | 0 | 0 | 0 | 0 | 0 | 0 | 0 | 0 | 0 |
| 17 | **0.53** | 0 | 0 | 0 | 0 | 0 | 0 | 0 | 1 | 0 |
| 18 | **0.73** | 0 | 0.565 | 0 | 0 | 0 | 0 | 0 | 1 | 0.733 |

Supplementary Table 4: Results for the generalised linear model (GLM) of the within-cluster variability in number of ignitions as a function of the annual gross primary production (GPP, $gCyr^{-1}$), maximum VPD (VPD, Pa), fractional grass cover (grass), fractional shrub cover (shrub), fractional tree cover tree), fractional cropland cover (crop), fractional pasture cover (pasture), road density (roads) and population density (popn). $R^2$ values for significant models are given in **bold.** Given that the sample size differs between clusters, the *t-values* were standardized but preserving the sign to reflect whether the variable had a positive or negative effect on number of ignitions. Thus, the standardised values vary between -1 and 1. Only significant *t-values* are shown.

| **cluster** | **r2** | **t values** | | | | | | | | |
|---|---|---|---|---|---|---|---|---|---|---|
| | | **tree** | **shrub** | **grass** | **popn** | **crop** | **pasture** | **roads** | **GPP** | **VPD** |
| 1 | **0.04** | 0 | 0 | 0 | 0 | 0 | 0 | 0 | 0 | 0 |
| 2 | **0.08** | 0 | -0.729 | 0 | 0 | 0 | 0 | -1 | -0.989 | 0 |
| 3 | **0.1** | -0.486 | 0 | 0.870 | 0 | 0 | 0.838 | 0 | 0 | 1 |
| 4 | **0.11** | 0.362 | 1 | -0.488 | 0 | 0.458 | 0.985 | -0.269 | -0.932 | 0.588 |
| 5 | **0.34** | 0 | -0.467 | -0.556 | 0 | 1 | 0 | 0 | 0 | 0 |
| 6 | **0.24** | 0 | -0.519 | -0.403 | 0 | 1 | 0 | -0.487 | 0 | -0.352 |
| 7 | **0.09** | 0 | 0.745 | 0.388 | 0 | 0 | -0.313 | 0.303732 | -1 | 0.474 |
| 8 | **0.31** | -0.611 | 0.528 | 0 | 0 | 0 | 0 | -0.976 | 0 | -1 |
| 9 | **0.24** | 0 | 0 | 0.473 | 0 | 0 | 0.417 | 0 | 0 | 1 |
| 10 | **0.17** | 0 | 0.833 | 0 | 0 | 0 | 0 | -0.273 | 0 | 1 |
| 11 | **0.37** | 0.959 | 0 | 1 | 0 | 0.325 | 0.564 | -0.231 | 0 | 0.175 |
| 12 | **0.16** | 0 | 0.453 | 0 | 0 | 0 | 1 | -0.628 | 0.602 | 0.456 |
| 13 | **0.43** | 0 | 0 | 0.175 | 0 | 0.133 | 0.684 | 0 | 1 | 0.785 |
| 14 | **0.35** | 0 | -0.298 | 0 | 0 | 0 | 1 | 0 | 0.473 | 0.445 |
| 15 | **0.62** | 0.343 | 0.443 | -0.396 | 0 | 0 | 0.292 | 0 | 1 | 0 |
| 16 | **0.09** | 0 | 0 | 0 | 0 | 1 | 0.711 | 0 | 0 | 0 |
| 17 | **0.11** | 0 | 0 | 0 | 0 | 0 | 0 | 0 | 0 | -1 |
| 18 | 0.52 | 0 | 0 | 0 | 0 | 0 | 0 | 0 | 0 | 0 |

Supplementary Table 5: Results for the generalised linear model (GLM) of the within-cluster variability in average fire size as a function of the annual gross primary production (GPP, $gCyr^{-1}$), maximum VPD (VPD, Pa), fractional grass cover (grass), fractional shrub cover (shrub), fractional tree cover tree), fractional cropland cover (crop), fractional pasture cover (pasture), road density (roads) and population density (popn). $R^2$ values for significant models are given in **bold.** Given that the sample size differs between clusters, the *t-values* were standardized but preserving the sign to reflect whether the variable had a positive or negative effect on number of ignitions. Thus, the standardised values vary between -1 and 1. Only significant *t-values* are shown.

| cluster | r2 | t values | | | | | | | | |
|---|---|---|---|---|---|---|---|---|---|---|
| | | tree | shrub | grass | popn | crop | pasture | roads | GPP | VPD |
| 1 | **0.33** | 0 | 0 | -0.796 | 0 | 0 | 0 | 0 | 0 | 1 |
| 2 | **0.27** | 0 | 0 | 0 | 0 | -0.797 | 0 | 0 | 0 | 1 |
| 3 | 0.14 | 0 | 0 | 0 | 0 | 0 | 0 | 0 | 0 | 0 |
| 4 | 0.17 | 0 | 0 | 0 | 0 | 0 | 0 | 0 | 0 | 0 |
| 5 | 0.32 | 0 | 0 | 0 | 0 | 0 | 0 | 0 | 0 | 0 |
| 6 | 0.34 | 0 | 0 | 0 | 0 | 0 | 0 | 0 | 0 | 0 |
| 7 | 0.2 | 0 | 0 | 0 | 0 | 0 | 0 | 0 | 0 | 0 |
| 8 | **0.37** | 0 | 0 | 0.503 | 0 | 0 | 0 | 0 | 0 | 1 |
| 9 | **0.12** | 0 | 0 | 0 | 0 | 0 | 0 | 0 | 0 | 1 |
| 10 | **0.08** | 0 | 0 | 0 | 0 | 0 | 0 | 0 | 0 | 0 |
| 11 | 0.31 | 0 | 0 | 0 | 0 | 0 | 0 | 0 | 0 | 0 |
| 12 | **0.24** | 0 | 0.734 | -0.356 | 0 | -0.733 | -0.551 | 0 | -0.950 | 1 |
| 13 | 0.52 | 0 | 0 | 0 | 0 | 0 | 0 | 0 | 0 | 0 |
| 14 | **0.29** | 0 | 0.995 | 0 | 0 | -1 | 0 | 0 | 0 | 0 |
| 15 | 0.46 | 0 | 0 | 0 | 0 | 0 | 0 | 0 | 0 | 0 |
| 16 | 0.46 | 0 | 0 | 0 | 0 | 0 | 0 | 0 | 0 | 0 |
| 17 | **0.2** | 0 | 0 | 0 | 0 | 0 | 0 | 0 | 0 | 0 |
| 18 | **0.91** | 0 | 0 | 0 | 0 | 0 | 0 | 0 | 0 | 0 |

Supplementary Table 6: Results for the generalised linear model (GLM) of the within-cluster variability in average fire speed as a function of the annual gross primary production (GPP, $gCyr^{-1}$), maximum VPD (VPD, Pa), fractional grass cover (grass), fractional shrub cover (shrub), fractional tree cover tree), fractional cropland cover (crop), fractional pasture cover (pasture), road density (roads) and population density (popn). $R^2$ values for significant models are given in **bold.** Given that the sample size differs between clusters, the *t-values* were standardized but preserving the sign to reflect whether the variable had a positive or negative effect on number of ignitions. Thus, the standardised values vary between -1 and 1. Only significant *t-values* are shown.

| **cluster** | **r2** | **t values** | | | | | | | | |
|---|---|---|---|---|---|---|---|---|---|---|
| | | **tree** | **shrub** | **grass** | **popn** | **crop** | **pasture** | **roads** | **GPP** | **VPD** |
| 1 | **0.11** | 0 | 0 | 0 | 0 | -0.740 | 0 | 0 | 0 | 1 |
| 2 | **0.14** | 0 | 1 | 0 | 0 | 0 | 0 | 0 | 0 | 0.768 |
| 3 | **0.1** | 0 | 0 | 0.937 | 0 | 0 | 0 | -1 | 0 | 0 |
| 4 | **0.12** | 0 | 0.784 | 0 | 0 | -1 | 0 | 0 | 0 | 0.989 |
| 5 | **0.22** | -0.283 | 0 | 0 | 0 | -1 | -0.489 | 0 | 0 | 0 |
| 6 | **0.32** | -0.430 | 0 | 0.335 | 0 | -1 | -0.283 | -0.430 | 0 | 0 |
| 7 | **0.14** | 0.976 | 0 | 0.707 | 0 | 0 | 0 | -0.444 | 0.388 | 1 |
| 8 | **0.04** | 0 | 0 | 0 | 0 | 0 | 0 | 0 | 0 | 0 |
| 9 | **0.12** | 0 | 0 | 0 | 0 | 0 | 0 | 0 | 0 | 1 |
| 10 | **0.04** | -0.934 | 0 | 0 | 0 | -1 | 0 | 0 | 0 | 0 |
| 11 | **0.25** | 0 | 1 | 0 | 0 | -0.419 | -0.802 | -0.478 | 0 | 0.863 |
| 12 | **0.14** | -0.313 | 0.434 | 0 | 0 | -0.302 | -0.279 | 0 | -1 | 0.535 |
| 13 | **0.36** | -0.623 | 0.442 | 0 | 0 | -1 | -0.815 | 0 | -0.429 | 0.979 |
| 14 | **0.22** | 0 | 1 | 0 | 0 | -0.560 | 0 | 0 | 0.527 | 0 |
| 15 | **0.31** | 0 | 0 | 0 | 0 | -1 | 0 | 0 | -0.726 | -0.625 |
| 16 | 0.38 | 0 | 0 | 0 | 0 | 0 | 0 | 0 | 0 | 0 |
| 17 | **0.20** | 0 | 0 | 0.946 | 0 | -0.681 | 0 | 0 | 0.848 | 1 |
| 18 | **0.61** | 0 | 0 | 0 | 0 | 0 | 0 | 0 | 0 | 0 |

Supplementary Table 7: Results for the generalised linear model (GLM) of the within-cluster variability in average fire duration as a function of the annual gross primary production (GPP, $gCyr^{-1}$), maximum VPD (VPD, Pa), fractional grass cover (grass), fractional shrub cover (shrub), fractional tree cover tree), fractional cropland cover (crop), fractional pasture cover (pasture), road density (roads) and population density (popn). $R^2$ values for significant models are given in **bold.** Given that the sample size differs between clusters, the *t-values* were standardized but preserving the sign to reflect whether the variable had a positive or negative effect on number of ignitions. Thus, the standardised values vary between -1 and 1. Only significant *t-values* are shown.

| cluster | r2 | t values | | | | | | | | |
|---|---|---|---|---|---|---|---|---|---|---|
| | | tree | shrub | grass | popn | crop | pasture | roads | GPP | VPD |
| 1 | **0.16** | 0 | 0 | 0 | 0 | 0 | 0 | 0 | 0 | 1 |
| 2 | **0.06** | 0 | 0 | 0 | 0 | 0 | 0 | 0 | 0 | 1 |
| 3 | **0.19** | 0.384 | 1 | 0 | 0 | 0 | 0 | -0.245 | -0.906 | 0.448 |
| 4 | **0.23** | 0.423 | 1 | -0.356 | 0 | -0.401 | -0.393 | 0 | 0 | 0.806 |
| 5 | **0.05** | 0.914 | 1 | 0 | 0 | 0 | -0.785 | 0 | 0 | 0 |
| 6 | **0.14** | 0 | -0.535 | 0 | 0 | -1 | 0 | 0 | 0 | 0 |
| 7 | **0.18** | 0.597 | 0 | 0 | 0 | 0 | 0 | -0.414 | 0.589 | 1 |
| 8 | **0.2** | 0 | 0 | 0 | 0 | 0 | 0 | 0 | 1 | 0 |
| 9 | **0.11** | -0.284 | 0 | 0 | 0 | 0 | 0 | 0.312 | 0 | 1 |
| 10 | **0.02** | -1 | 0 | 0 | 0 | 0 | 0 | 0 | 0 | 0 |
| 11 | **0.19** | 0 | 0.760 | -0.939 | 0 | -0.652 | -1 | -0.457 | 0.389 | 0 |
| 12 | **0.04** | 0 | 1 | 0 | 0 | 0 | -0.590 | 0 | -0.434 | 0 |
| 13 | **0.16** | 0 | 0.498 | 0 | 0 | -0.576 | -1 | 0 | 0.440 | 0.225 |
| 14 | **0.11** | 0 | 1 | 0 | 0 | 0 | 0 | 0 | 0.673 | 0 |
| 15 | **0.09** | 0 | 0 | 0 | 0 | -0.839 | 0 | 0 | 0 | -1 |
| 16 | **0.34** | 0 | -0.257 | -0.736 | 0 | -0.400 | 0 | -0.560 | 1 | 0.307 |
| 17 | **0.19** | 0 | 0 | 0 | 0 | 0 | 0 | 0 | 1 | 0 |
| 18 | 0.47 | 0 | 0 | 0 | 0 | 0 | 0 | 0 | 0 | 0 |

Supplementary Table 8: Results for the generalised linear model (GLM) of the within-cluster variability in carbon emissions as a function of the annual gross primary production (GPP, $gCyr^{-1}$), maximum VPD (VPD, Pa), fractional grass cover (grass), fractional shrub cover (shrub), fractional tree cover tree), fractional cropland cover (crop), fractional pasture cover (pasture), road density (roads) and population density (popn). $R^2$ values for significant models are given in **bold.** Given that the sample size differs between clusters, the *t-values* were standardized but preserving the sign to reflect whether the variable had a positive or negative effect on number of ignitions. Thus, the standardised values vary between -1 and 1. Only significant *t-values* are shown.

| **cluster** | **r2** | **t values** | | | | | | | | |
|---|---|---|---|---|---|---|---|---|---|---|
| | | **tree** | **shrub** | **grass** | **popn** | **crop** | **pasture** | **roads** | **GPP** | **VPD** |
| 1 | **0.58** | 0 | 0.818 | -0.720 | 0 | 0 | 0 | 0 | 1 | 0.913 |
| 2 | **0.19** | 0 | 0 | 0 | 0 | 0 | 0 | 0 | 0 | 0 |
| 3 | **0.26** | 0 | 0.192 | 0.243 | 0 | 0 | 0.171 | -0.291 | -0.255 | 1 |
| 4 | **0.30** | 0 | 0.404 | 0 | 0 | 0 | 0 | 0 | 0 | 1 |
| 5 | **0.22** | 0 | 0 | 0 | 0 | 0.480 | 0 | 0 | 1 | 0.213 |
| 6 | **0.25** | -0.227 | -0.482 | 0 | 0 | 0 | -0.164 | 0 | 1 | 0 |
| 7 | **0.32** | 0.343 | 0.338 | 0.177 | 0 | 0 | 0 | 0 | 0 | 1 |
| 8 | **0.39** | 0.410 | 1 | 0 | 0 | 0 | 0 | -0.368 | 0.796 | 0.418 |
| 9 | **0.41** | -0.250 | 0.195 | 0.165 | 0 | 0 | 0.156 | 0 | 0 | 1 |
| 10 | **0.28** | 0 | 0.832 | 0 | 0 | 0 | 0 | 0 | 0 | 1 |
| 11 | **0.49** | 1 | 0.107 | 0.962 | 0 | 0 | 0.357 | -0.118 | 0.381 | 0.411 |
| 12 | **0.28** | 0 | 0.458 | 0 | 0 | 0 | 0.325 | -0.289 | 1 | 0.647 |
| 13 | **0.71** | 0.030 | 0.157 | 0.151 | 0 | 0 | 0.110 | -0.060 | 1 | 0.637 |
| 14 | **0.69** | 0 | 0 | 0 | 0 | 0 | 0.385 | 0 | 1 | 0.592 |
| 15 | **0.9** | 0.125 | 0.266 | -0.10 | 0 | -0.161 | 0 | 0.073 | 1 | 0 |
| 16 | 0.68 | 0 | 0 | 0 | 0 | 0 | 0 | 0 | 0 | 0 |
| 17 | **0.68** | 0 | 0.194 | 0 | 0 | 0.246 | 0.224 | -0.166 | 1 | 0.565 |
| 18 | **0.73** | 0.566 | 0 | 0 | 0 | 0 | 0 | 0 | 1 | 0 |

Supplementary Table 9: Results for the generalised linear model (GLM) of the within-cluster variability in fire seasonal concentration as a function of the annual gross primary production (GPP, $gCyr^{-1}$), maximum VPD (VPD, Pa), fractional grass cover (grass), fractional shrub cover (shrub), fractional tree cover tree), fractional cropland cover (crop), fractional pasture cover (pasture), road density (roads) and population density (popn). $R^2$ values for significant models are given in **bold.** Given that the sample size differs between clusters, the *t-values* were standardized but preserving the sign to reflect whether the variable had a positive or negative effect on number of ignitions. Thus, the standardised values vary between -1 and 1. Only significant *t-values* are shown.

| **cluster** | **r2** | **t values** | | | | | | | | |
|---|---|---|---|---|---|---|---|---|---|---|
| | | **tree** | **shrub** | **grass** | **popn** | **crop** | **pasture** | **roads** | **GPP** | **VPD** |
| 1 | **0.07** | 0 | 0 | 0 | 0 | -0.787 | 1 | 0.864 | 0 | 0 |
| 2 | **0.07** | 0 | 0 | 0 | 0 | 0 | 0 | 0.814 | 1 | -0.822 |
| 3 | **0.22** | 0.853 | 0.539 | 0 | 0 | -0.649 | 0 | 0.348 | -0.329 | -1 |
| 4 | **0.36** | 0.362 | 0 | 0.190 | -0.133 | -1 | 0 | 0.211 | 0 | -0.415 |
| 5 | **0.05** | 0 | 0 | 0 | 0 | -0.579 | 0 | 0 | -1 | -0.499 |
| 6 | **0.03** | 0 | 0 | 0 | 0 | 0 | 0 | 0 | -1 | 0 |
| 7 | **0.05** | 0 | 0 | 0 | 0 | -0.901 | 0 | -1 | -0.678 | 0 |
| 8 | **0.18** | 0 | 0 | 0.533 | 0 | -1 | 0 | -0.785 | -0.497 | 0 |
| 9 | **0.03** | 0.807 | 0 | 0 | 0 | 0 | 0 | 0 | 0.672 | 1 |
| 10 | **0.11** | 0.379 | 1 | -0.955 | 0 | 0 | 0 | 0 | 0 | 0 |
| 11 | **0.13** | 1 | 0 | 0.701 | 0 | 0 | 0.548 | 0 | 0 | 0.853 |
| 12 | **0.05** | 1 | 0.457 | -0.436 | 0.938 | 0 | 0.969 | 0 | 0 | 0 |
| 13 | **0.03** | 0 | 0 | 0 | 0 | 0 | 0.846 | 0 | 0 | 1 |
| 14 | **0.03** | 0 | 0 | 0 | 0 | 0 | 0.972 | 0 | -1 | -0.824 |
| 15 | **0.06** | 0 | 0 | 0 | 0.715 | 0 | 0 | -0.987 | 0 | -1 |
| 16 | **0.04** | 0 | 0 | 0 | 0 | 0.523 | 0 | 0 | -1 | -0.775 |
| 17 | **0.05** | 0 | 0 | 0 | 0 | 0 | 0 | 0 | -1 | -0.904 |
| 18 | **0.59** | 0 | 0 | 0 | 0 | 0 | -1 | 0 | 0 | 0 |
